# Robust transcriptional indicators of plant immune cell death revealed by spatio-temporal transcriptome analyses

**DOI:** 10.1101/2021.10.06.463031

**Authors:** Jose Salguero-Linares, Irene Serrano, Nerea Ruiz-Solani, Marta Salas-Gómez, Ujjal Jyoti Phukan, Victor Manuel González, Martí Bernardo-Faura, Marc Valls, David Rengel, Nuria S. Coll

**Affiliations:** Centre for Research in Agricultural Genomics (CRAG), CSIC-IRTA-UAB-UB, Campus UAB, Bellaterra, Barcelona, 08193, Spain; LIPM, Universite de Toulouse, INRA, CNRS, 84195 Castanet-Tolosan, France; INRAE, GeT-PlaGe, Genotoul, 31326 Castanet-Tolosan, France (doi: 10.15454/1.5572370921303193E12); Department of Genetics, Universitat de Barcelona, 08028 Barcelona, Spain; Consejo Superior de Investigaciones Científicas (CSIC), Barcelona, Spain

**Keywords:** *Arabidopsis thaliana*, Cell Death Indicator, Effector-Triggered Immunity, Hypersensitive Response, Immune Cell Death, Pattern-Triggered Immunity, Plant Immunity, *Pseudomonas syringae*

## Abstract

Recognition of a pathogen by the plant immune system often triggers a form of regulated cell death traditionally known as the hypersensitive response. This type of immune cell death occurs precisely at the site of pathogen recognition, and it is restricted to a few cells. Extensive research has shed light into how plant immune receptors are mechanistically activated. However, a central key question remains largely unresolved: how does cell death zonation take place and what are the mechanisms that underpin this phenomenon? As a consequence, *bona fide* transcriptional indicators of immune cell death are lacking, which prevents gaining a deeper insight of its mechanisms before cell death becomes macroscopic and precludes any early or live observation. We addressed this question using the paradigmatic *Arabidopsis thaliana*–*Pseudomonas syringae* pathosystem, by performing a spatio-temporally resolved gene expression analysis that compared infected cells that will undergo immune cell death upon pathogen recognition *vs* by-stander cells that will stay alive and activate immunity. Our data revealed unique and time-dependent differences in the repertoire of differentially expressed genes, expression profiles and biological processes derived from tissue undergoing immune cell death and that of its surroundings. Further, we generated a pipeline based on concatenated pairwise comparisons between time, zone and treatment that enabled us to define 13 robust transcriptional immune cell death markers. Among these genes, the promoter of an uncharacterized *AAA-ATPase* has been used to obtain a fluorescent reporter transgenic line, which displays a strong spatio-temporally resolved signal specifically in cells that will later undergo pathogen-triggered cell death. In sum, this valuable set of genes can be used to define those cells that are destined to die upon infection with immune cell death-triggering bacteria, opening new avenues for specific and/or high-throughput techniques to study immune cell death processes at a single-cell level.

## INTRODUCTION

Plants are rich sources of nutrients for pathogens with contrasting lifestyles (1). As opposed to animals, plants do not possess a circulatory system with mobile cells specialized in pathogen defense (2). Since their cells are fixed by their cell walls, plants rely on each cell’s autonomous immunity and on systemic signals emanating from infection sites to distal cells to prime the plant for future pathogen encounters (3). Moreover, instead of a somatic adaptive immune system that produces antigen receptors on demand, plant cells are equipped with extracellular pattern-recognition receptors (PRRs) and intracellular nucleotide-binding leucine-rich repeat immune receptors (NLRs) that recognize microbe-associated microbial patterns (MAMPs) and pathogen effectors required for virulence, respectively (4). PRR activation brings about a broad defense response named pattern-triggered immunity (PTI), while NLR activation triggers a potentiated and prolonged immune response named effector-triggered immunity (ETI) that reinforce defense outputs observed during PTI (5, 6). ETI often culminates in a macroscopic localized cell death at the attempted pathogen ingress site known as hypersensitive response (HR)-cell death or immune-related cell death (7–9).

Regulated cell death has a crucial role in both animals and plant immune responses. Extensive research in the animal field supports the notion that the immune system is highly dependent on cell death for a robust and tightly controlled immune response to occur (10, 11). In plants, our knowledge about the biochemical and genetic pathways regulating cell death, particularly in the context of immunity, is still very limited. As an attempt to shed light into how immune cell death is orchestrated in plants, most efforts have been directed towards understanding how NLRs are mechanistically activated, as well as identifying molecular components upstream or downstream of NLRs that are required for HR to occur (12–14).

Plant NLRs can be broadly classified into TNLs and CNLs based on their domain composition: TNLs contain a Toll/Interleukin-1 Receptor (TIR), whereas CNLs harbor a coiled-coiled domain at their N-terminal end (15). Groundbreaking research has shown that, in plants, activation of certain NLRs by pathogen perception is mediated by oligomerization, which ultimately will result in cell death and immunity (13, 14). Oligomerized forms of CNLs can potentially form pores at the plasma membrane (16, 17). Some TNLs, in turn, can oligomerize upon activation to reconstitute a holoenzyme (NAD hydrolase) that triggers cell death by a mechanism that is not fully elucidated (18–20). Further, a subset of NLRs known as “helper NLRs” (or RNLs) are part of networks that function downstream pathogen-sensing or “sensor” NLRs and some of them have been shown to oligomerize and form calcium channels at the plasma membrane (17, 21, 22). However, it remains unclear how oligomerization translates to immune signaling and immune cell death.

In the context of signaling downstream NLR activation or ETI, large-scale transcriptional studies have highlighted the importance of phytohormone networks for high-amplitude transcriptional reprogramming to mount a fast and efficient response (23). Comparisons between host transcriptional responses elicited by PTI and ETI suggest minor qualitative differences in the repertoire of genes differentially expressed (23, 24). These studies also support the recently evidenced assumption that ETI and PTI share immune signaling components (5, 6, 25, 26). However, a central key question remains unexplored: which early transcriptional signatures differentiate cells that recognize the pathogen and will undergo ETI immune cell death from by-stander cells that will remain alive and will activate defenses to fight the pathogen? In recent literature, a few studies underscore the importance of zonation during immune cell death (27–29). At the hormonal level, it has been shown that salicylic acid (SA) plays a major role at pathogen-inoculated spots that will later undergo immune cell death, while the jasmonic acid (JA) signaling pathway is activated in the cells surrounding the central SA-active cells (29). Furthermore, precision transcriptomics during the immune response elicited by the potato Ny-1 gene against potato virus Y (PVY) revealed the importance of SA accumulation and genes involved in the generation of reactive oxygen species (ROS) for efficient confinement of macroscopic cell death lesions caused by PVY (27). The cell wall polymer lignin has also been shown to participate in immune cell death zonation, by forming a physical barrier around the infection site upon pathogen recognition that presumably contributes to confine the invading agents and restricts colonization (30). A transcriptional meta-analysis of developmental vs immune cell death in plants could only reveal robust indicators for developmental cell death but not of immune cell death (7). We realized that the limitation of previous large-scale transcriptomic analysis lacked the spatial dimension of immune cell death (23, 31), as dying cells were not compared to by-stander cells, and the focus was not placed on identifying specific cell death markers, but rather bulk-analyzing the ETI response at the inoculated area.

A systematic gene expression analysis of the zonation of immune cell death overtime would help understanding the process of immune cell death at the molecular level and importantly, would allow defining *bona fide* transcriptional markers of the process. With this purpose, we generated RNA-sequencing (RNA-seq) data to systematically analyze and compare the transcriptional programs taking place at the zone of inoculation/pathogen recognition that will undergo immune cell death vs the surrounding area that will stay alive and activate immunity. We show unique and time-dependent differences in the repertoire of differentially expressed genes (DEGs) and expression profiles derived from tissue undergoing immune cell death and that of its surroundings. Furthermore, we generated a pipeline based on pairwise comparisons between time, zone and treatment that enabled us to define 13 robust transcriptional immune cell death markers and a fluorescent transgenic reporter line. These valuable set of genes can be used to define those cells that are destined to die upon pathogen recognition before the onset of cell death becomes macroscopically visible, opening new horizons to study the processes therein by live, cell-specific and/or high-throughput techniques.

## RESULTS

### Zonally dissected Arabidopsis transcriptomes upon *Pto AvrRpm1* infection reveal unique spatio-temporal gene expression

In our experiments we used the paradigmatic interaction between *Arabidopsis thaliana* Col-0 (hereafter Arabidopsis) and the bacterial pathogen *Pseudomonas syringae* pathovar tomato (*Pto*) carrying the effector *AvrRpm1* (hereafter *Pto AvrRpm1)*, which triggers restricted immune cell death at the site of inoculation upon recognition by the CNL RPM1 (RESISTANCE TO PSEUDOMONAS SYRINGAE PV MACULICOLA 1) (32). In order to zonally dissect immune cell death and its surrounding, we syringe-infiltrated a limited area (roughly 3-4 mm) at the side edge of Arabidopsis leaves with either a mock solution or *Pto AvrRpm1.* Collected tissue from this area was designated as the “IN” zone. To ensure proper separation between IN and OUT zones, a buffer zone expanding 1 mm next to the IN area was discarded, and a parallel region expanding 1 to 2 mm towards de vein was designated as “OUT” (Figure 1a). We collected tissue at 0, 1-, 2-, 4- and 6-hours post-inoculation (hpi), extracted RNA and assessed transcript abundance by RNA-seq. Under these conditions, macroscopic cell death started appearing at 4 hpi in the *Pto AvrRpm1*-inoculated samples, as visualized by trypan blue staining (Figure 1b). As expected, this cell death is concomitant with a dramatic drop in photosynthetic efficiency of photosystem II (Fv/Fm ratio) and electron transport rate (ETR) at the IN area (Figure 1c) (33).

**Figure 1.**
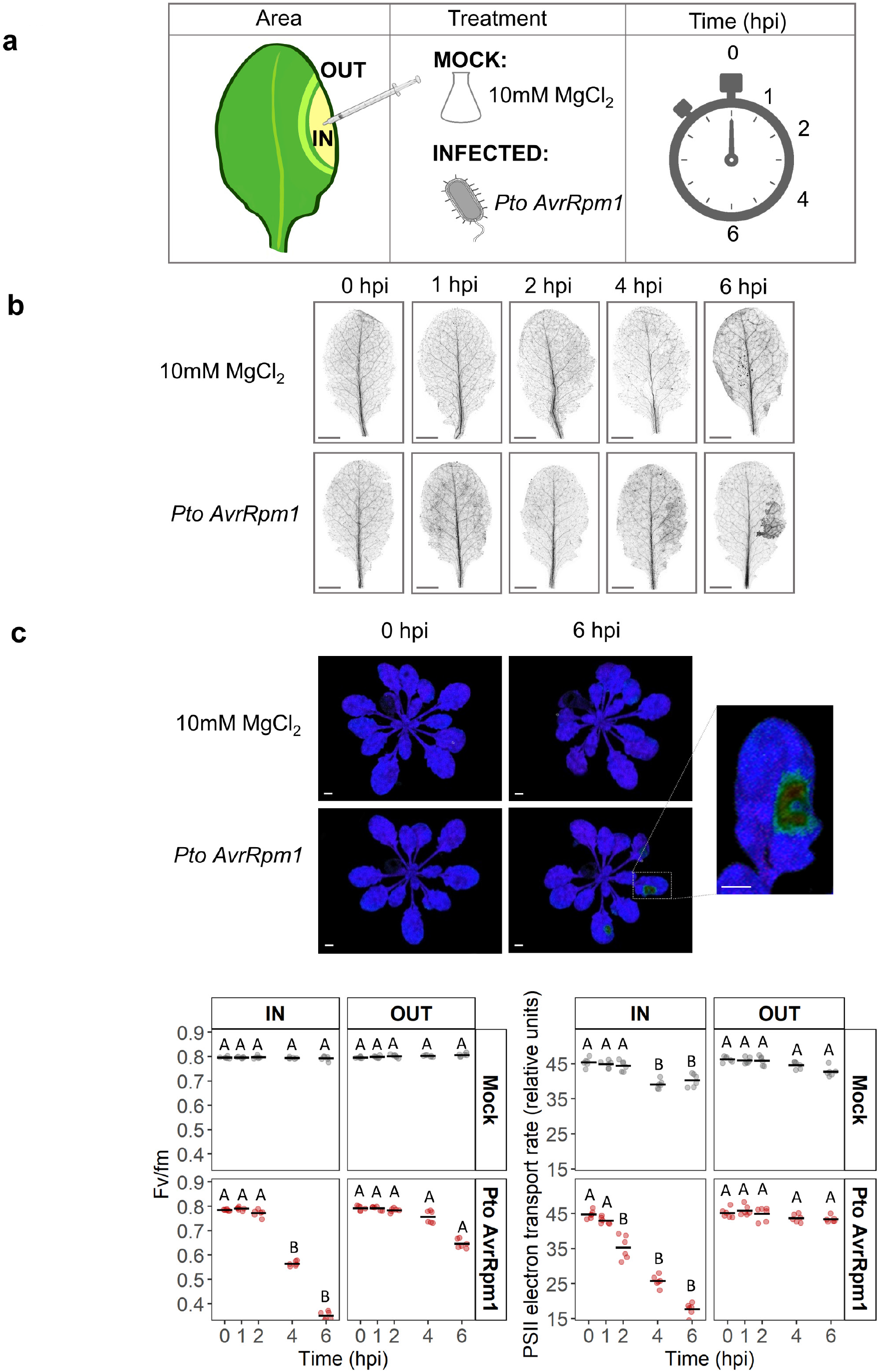
Immune cell death in plants can be spatio-temporally dissected. **(a)** Experimental design of the study. A limited area (3-4 mm) at the side edge of four-week-old Arabidopsis thaliana Col-0 leaves was syringe-infiltrated with either *Pto AvrRpm1* at 2.5*10^7^ cfu/ml (INFECTED) or a 10 mM MgCl_2_ solution (MOCK) and samples were collected at 5 different time points after infection: 0, 1, 2, 4 and 6 hpi. Upon infiltration, the edge of the infiltrated area was marked, and the total area infiltrated designated as “IN”. A 1 mm buffer zone right next to the IN zone ensured proper separation between the IN and “OUT” area, which was the parallel region that expanded from the edge of the buffer zone to 1-2 mm towards the vein. Three biological replicates per area, treatment and time point were collected and subjected for RNA-seq analysis. **(b)** Analysis of macroscopic cell death upon infection with either *Pto AvrRpm1* or 10 mM MgCl_2_ solution. Leaves were infected as described in (a) and subsequently stained with trypan blue. Scale bar 3 mm **(c)** Representative images of mock or *Pto AvrRm1*-treated plants subjected to pulse-amplitude modulated (PAM) chlorophyll fluorescence measurement to monitor photosynthesis. Scale bar 3 mm. Photosynthetic efficiency (Fv/Fm ratio) and electron transport rate (ETR) were measured in the infiltrated area (IN) and the neighboring tissue (OUT). Measurements were taken at 0, 1, 2, 4 and 6 hpi. Results are representative of 6 different measurements of each tissue area from 6 different plants. Letters indicate statistically significant differences in either Fv/Fm ratio or ETR values following a two-way ANOVA with Tukeýs HSD test (α = 0.05). Exact p values are provided in **Table S2**.

To determine whether the obtained RNA-seq data complied with our working hypothesis of spatio-temporal gene expression regulation we performed a Principal Component Analysis (PCA) (Figure S1). We observed that at the IN area, *Pto AvrRpm1*-treated samples separated from their mock controls from 2 hpi onwards. At the OUT area, however, only *Pto AvrRpm1*-treated samples at 4 and 6 hpi separated from mock controls. Overall, the PCA confirms that the biggest changes in gene expression are produced at IN, particularly at 4 and 6 hpi, whereas at OUT there is a subtler modulation that is most pronounced at 4 hpi.

Next, we identified differentially expressed genes (DEGs) between bacteria and mock-inoculated samples (DEGs; false discovery rate (FDR) < 0.05 and |log2FC| > 2), thereby characterizing the transcriptional changes occurring at each tissue area at every time point. We found a total of 5,495 DEGs at the IN zone and 1,785 at the OUT zone (Figure 2a). Enrichment of Gene Ontology (GO) terms was examined in every group of DEGs at each specific time point (Figure S2). Upregulated genes at the IN area were enriched in immunity- and phytohormone-associated processes (Figure S2a). Immunity-related GO terms associated with PTI and ETI appeared at initial stages of infection (1 and 2 hpi), while at later stages (from 2 hpi onwards) an enrichment in genes involved in more general defense and abiotic stress responses could be observed (Figure S2a). Regarding phytohormone-related processes, we observed an enrichment in SA-related GO terms from 1 hpi onwards, confirming the importance of SA at the IN area (34). In contrast, GO terms associated with JA were particularly overrepresented at later time points (4 and 6 hpi), in accordance with previous findings demonstrating that SA can activate JA signaling through a non-canonical pathway promoting ETI (35). GO terms related to other defense/stress-related phytohormones such as ethylene (ET) and abscisic acid (ABA), were also enriched at 4 and 6 hpi (Figure S2a).

**Figure 2.**
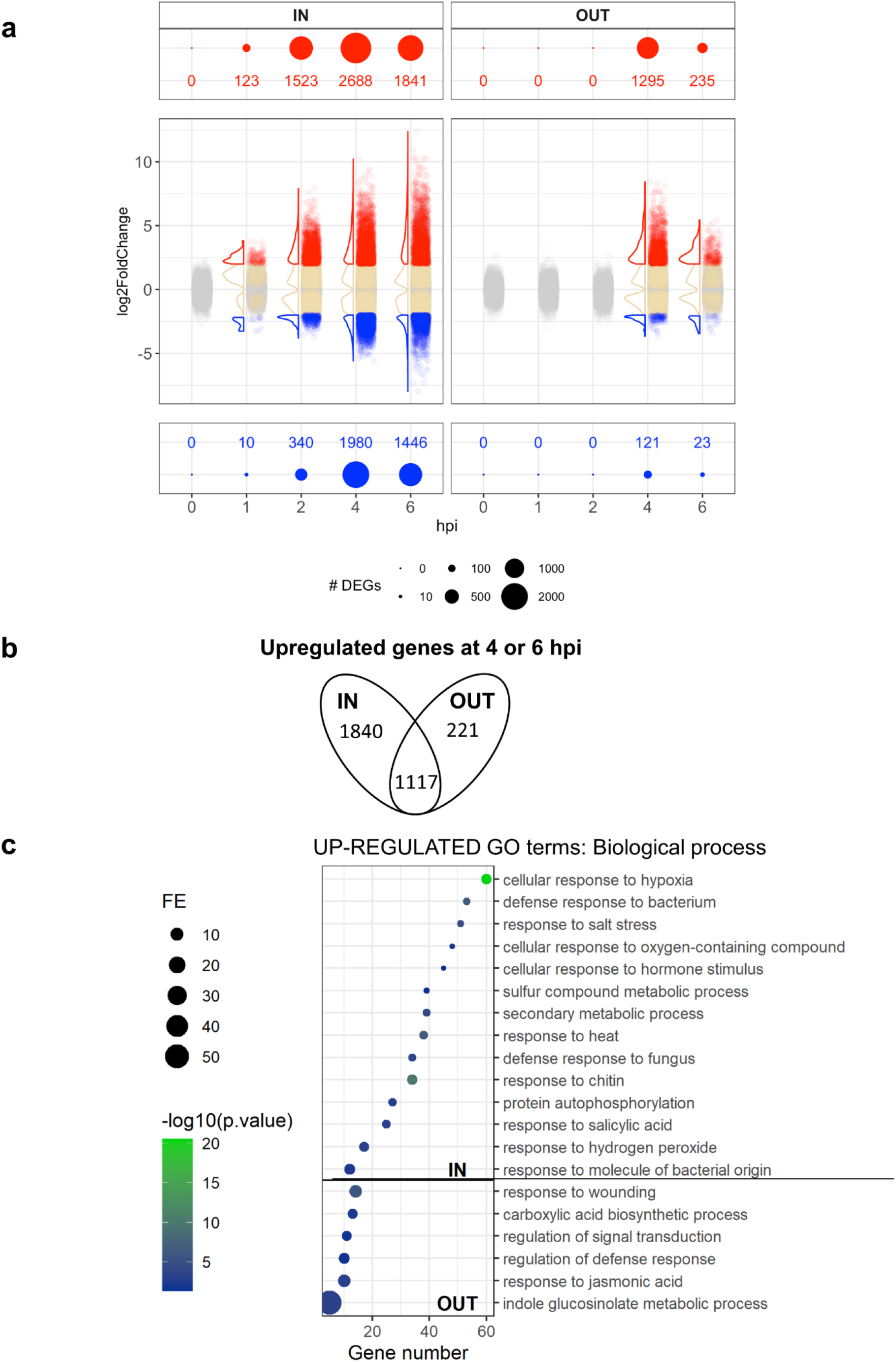
Spatio-temporal dynamics of the transcriptome reveal time and zone-dependent gene expression signatures upon infection. **(a)** Differentially expressed genes (FDR < 0.05 and |log_2_FC| > 2) in *Pto AvrRpm1*-infected plants compared to mock-treated plants at each time point at the IN (left) and OUT (right) areas. Red dots denote upregulated genes and blue dots indicate downregulated genes. **(b-c)** Genes exclusively upregulated (FDR < 0.05 and log_2_FC>2) at either IN or OUT areas of infection at 4 and 6 hpi. **(b)** Venn diagrams showing sizes of gene sets that are upregulated upon bacterial infection at 4 or 6 hpi at either IN, OUT or both areas. **(c)** GO terms representing enriched biological processes derived from genes exclusively upregulated at either IN or OUT areas. The most specific term from each family term provided by PANTHER was plotted along with their corresponding gene number, fold enrichment (FE) and FDR (Bonferroni Correction for multiple testing) represented as log_10_. Only GO Terms with a FE above 2 and FDR below 0.05 were plotted.

Among downregulated genes at the IN zone, an enrichment in GO terms related to photosynthesis and chloroplast biology occurred at late time points (4 and 6 hpi) (Figure S2b). This correlates with the drop in photosynthetic efficiency shown in Figure 1c, which is part of the defense/yield trade-off to derive resources to immune responses and shut down production of sugars and nutrients, as they might serve as a source for pathogen survival and multiplication (36).

Strikingly, at the OUT area we only observed differential expression at late time points (4 and 6 hpi), with an overall reduction in the number of DEGs compared to the IN area (Figure 2a). Upregulated genes were enriched in GO terms associated with hormonal regulation, particularly to the JA signaling pathway (Figure S2c). Downregulated genes at the OUT area did not show any particular enriched GO term, possibly due to the low number of genes.

To identify genes exclusively upregulated at either the IN or OUT areas we first generated Venn diagrams representing the sizes of gene sets induced at each time point upon infection (Figure S3). This analysis confirmed that upregulation at both IN and OUT mainly occurs at 4 or 6 hpi (Figure S3) and revealed genes exclusively upregulated at the IN and OUT areas at these time points (Figure 2b). Specifically, we found a total of 1,840 genes being upregulated exclusively at IN, 1,117 genes upregulated at both IN and OUT and 221 genes being exclusively upregulated at OUT (Figure 2b). Among the overrepresented GO terms found in genes exclusive for the IN area were “defense response to bacterium”, “response to molecule of bacterial origin” or “response to salicylic acid”. We also found various GO terms associated to responses to several other stresses such as salt, oxygen-containing compounds, sulfur compounds, heat and hydrogen peroxide (Figure 2c), which is not surprising, considering that the tissue is undergoing cell death. In contrast, overrepresented GO terms in genes exclusively upregulated at the OUT area included “regulation of defense response” and, interestingly, “response to wounding” and “response to jasmonic acid” (Figure 3b). These JA-related genes follow a very distinct expression pattern, with an early peak at 1 hpi both at the IN and OUT areas, and a second peak at 4 hpi of higher intensity in the OUT zone (Figure S4, **Table S5**). Although further experimental validation would be required, these data reveal expression patterns of a set of genes that could potentially be used as OUT markers along with previously reported markers such as VSP1 (29, 37).

**Figure 3.**
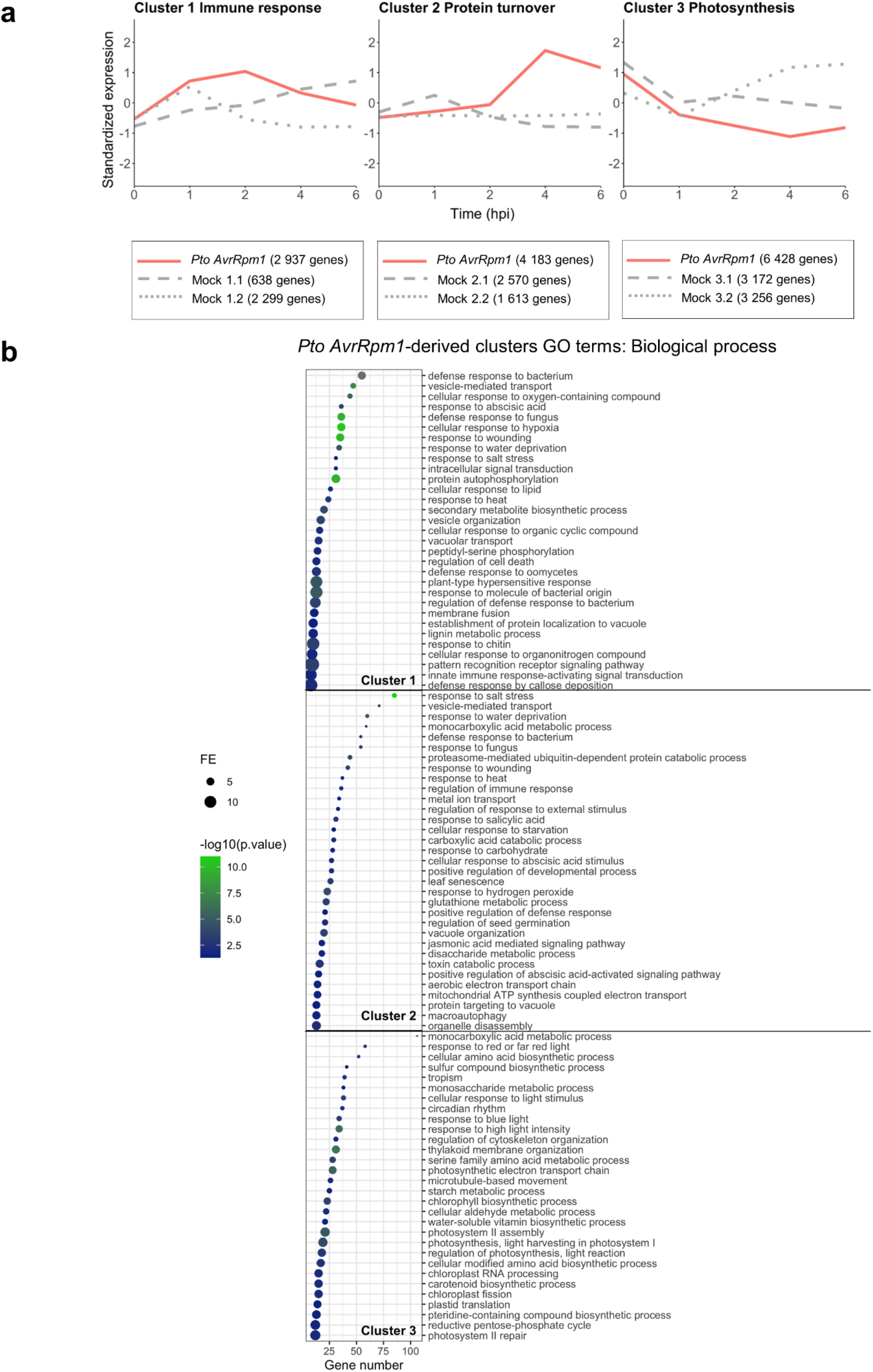
Gene expression profile clustering at the IN area reveals three distinctive expression patterns related to immunity, protein turnover and photosynthesis. **(a)** Non-overlapping clusters derived from *Pto AvrRpm1-* and mock-treated plants at the IN area. The trajectory that defines the overall expression profile of each cluster is shown in red for *Pto AvrRpm1*-treated plants. Genes derived from *Pto AvrRpm1-*treated samples were re-clustered for mock-treated samples and their trajectories are represented in grey. Since the expression profile of these genes in mock-treated samples was very distinct among the overall number of genes, they were divided into two sub-clusters represented either in dotted or dashed grey lines. The number of genes that constitute each cluster is indicated. **(b)** GO terms representing enriched biological processes derived from each cluster in *Pto AvrRpm1*-treated plants. GO term enrichment analysis was performed on those genes that had a membership score value (MSV) above or equal to 0.7 (See Materials and Methods). The most specific term from each family provided by PANTHER was plotted along with their corresponding gene number, fold enrichment (FE) and FDR (Bonferroni Correction for multiple testing) represented as log_10_. Only GO Terms with a FE above 2 and FDR below 0.05 were plotted. Enriched GO terms from cluster 1 (2,937 genes; MSV > 0.7 → 1069 genes), cluster 2 (4,183 genes; MSV > 0.7 → 2613 genes) and cluster 3 (6,428 genes; MSV > 0.7 → 4885 genes) in *Pto AvrRpm1*-treated plants were predominantly linked to processes related to immunity, protein turnover and photosynthesis, respectively.

### Clustering of gene expression profiles reveals distinct expression patterns at the IN and OUT areas over time

Next, we set out to determine whether genes at the IN and OUT areas followed specific expression patterns and if particular biological processes were associated to those patterns. For this, we first analyzed gene expression profiles using Fuzzy c-means, a soft partitioning algorithm which offers robust clustering with regards to noise by variation of a fuzzification parameter that limits the contribution of ill-behaved profiles to the clustering process (38, 39). Based on this, we could define three and five distinct and non-overlapping clusters for *Pto AvrRpm1*-treated samples in the IN (Figure 3a) and OUT (Figure 4a **and** Figure S7) areas, respectively. Genes within each cluster were subsequently re-clustered in mock-treated samples, producing two distinct sub-clusters (Figures 3a and 4a). This procedure provided a more detailed overview of the differences and similarities of trajectories between treatments over time and reflected the well-documented wound response that takes place in mock-treated tissue (23, 28, 40)

**Figure 4.**
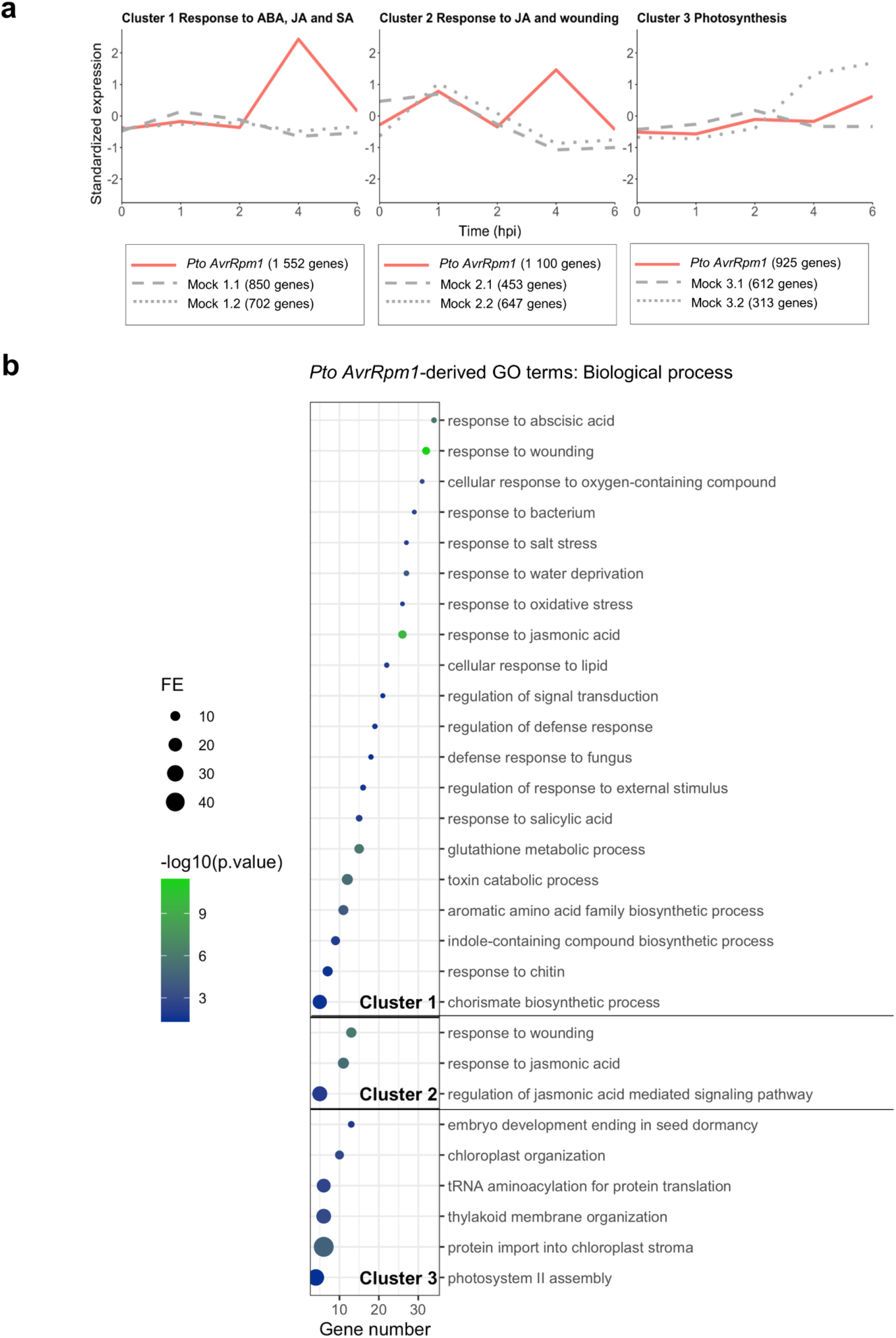
Gene expression profile clustering at the OUT area reveals the importance of phytohormone regulation around the infection site. **(a)** Non-overlapping clusters derived from *Pto AvrRpm1-* and mock-treated plants at the OUT area of infection. The trajectory that defines the overall expression profile of each cluster is shown in red for *Pto AvrRpm1*-treated plants. Genes derived from *Pto AvrRpm1-*treated samples were re-clustered for mock-treated samples and their trajectories are represented in grey. Since the expression profile of these genes in mock-treated samples was very distinct among the overall number of genes, they were divided into two sub-clusters represented either in dotted or dashed grey lines. The number of genes that constitute each cluster is indicated. **(b)** GO terms representing enriched biological processes derived from each cluster in *Pto* (*AvrRpm1*)-treated plants. GO term enrichment analysis was performed on those genes that had a membership score value (MSV) above or equal to 0.7. The most specific term from each family provided by PANTHER was plotted along with their corresponding gene number, fold enrichment (FE) and FDR (Bonferroni Correction for multiple testing) represented as log_10_. Only GO Terms with a FE above 2 and FDR below 0.05 were plotted. Enriched GO terms from cluster 1 (1,552 genes; MS > 0.7 → 747 genes) and 2 (1,100 genes; MS > 0.7 → 184) suggest the importance of processes related to hormonal regulation in by-stander cells, whereas genes comprising cluster 3 (925 genes; MS > 0.7 → 181 genes) infer that photosynthesis and rearrangements in the chloroplast occur similarly compared to mock-treated samples at the OUT area.

At the IN area of infection, cluster 1 exhibited a pattern of upregulation from 0 to 2 hpi and mild downregulation from 2 to 6 hpi. We dubbed this cluster “Immune response” as genes near to its centroid (see Materials and Methods) are mainly associated with immune-related GO terms (Figure 3b). Genes in this cluster followed two distinct trajectories in the mock-treated samples: while mock sub-cluster 1.1 showed a steady increase throughout the experiment, mock sub-cluster 1.2 exhibited a typical wounding immune-related response common with infected samples, peaking at 1 h and rapidly returning to steady-state levels (41). The different behavioral pattern of genes observed from 1 to 2 hpi in infected IN samples with respect to the mock treatment reveals a specific response to bacterial infection taking place specifically at the site of infection (Figure 3a **and** Figure S5).

Cluster 2-IN includes genes with a sharp increase of expression at 4 hpi (Figure 3a). We termed this cluster “Protein turnover” as genes following that trajectory are to a certain extent involved in protein degradation processes (autophagy, protein targeting to the vacuole, proteasome mediated degradation) taking place in response to infection (Figure 3b). Sub-clusters from mock-treated samples predominantly followed a similar steady trajectory throughout the experiment, which points to an infection-specific effect of upregulation on protein turnover due to infection at the IN area (Figure 3 **and** Figure S5).

Cluster 3-IN exhibits an expression pattern of steady downregulation from 0 to 4 hpi, followed by a slight recovery of expression from 4 to 6 hpi (Figure 3a). We designated cluster 3 as “Photosynthesis” as it includes mostly genes belonging to GO terms related to this process (Figure 3b). In this case, mock-treated samples sub-cluster into two distinct patterns of expression: sub-cluster 3.1 follows a similar pattern as infected samples, while sub-cluster 3.2 shows a transient decrease of expression at 1 h followed by a recovery phase from 2 to 6 hpi (Figure 3a). We conclude that only certain components of the photosynthetic machinery are specifically affected by the pathogen treatment (“photosynthetic electron transport chain”, and “thylakoid membrane organization”), whereas other aspects of photosynthesis behave similarly overtime regardless of infection (“photosynthesis, light harvesting in photosystem” and “regulation of photosynthesis”) (Figure 3b **and** Figure S5).

At the OUT area of infection, cluster 1 includes genes that display a sharp peak of expression at 4 hpi (Figure 4a). From this cluster, genes near the centroid belong to GO terms associated with metabolism, hormonal regulation, and wounding response, among others. Interestingly, JA- and SA-responsive genes, which are known to act antagonistically and cooperatively during ETI (29, 35), seem to be highly enriched in the OUT area, consistent with previous studies that considered the spatiotemporal dimension of cell death (27) (Figure 4b). The behavior of the genes that comprise cluster 1-OUT in infected samples is remarkably different in mock-treated samples (Figure 4a). Genes comprising the mock-derived sub-clusters follow a similar trend of steady expression throughout the time course of the experiment, suggesting that the peak of high expression is a specific response to the bacterial infection in the surrounding area (Figure 4a **and** Figure S6).

Cluster 2-OUT in *Pto AvrRpm1*-treated samples follows an expression pattern with two sharp upregulation peaks at 1 and 4 hpi (Figure 4a). These trajectories are followed by genes associated with JA-related processes and wounding, and is a very specific pattern exclusively found at the OUT zone (Figure 2b). Interestingly, both mock sub-clusters in this category, display a peak of upregulation at 1 hpi but not at 4 hpi. The early peak at 1 hpi shared between mock and infected samples could account for a wounding response elicited early at the area surrounding the syringe-infiltrated area. In contrast, the second peak at 4 hpi appears as a late response that occurs specifically at the tissues surrounding the pathogen inoculation area (Figure 4a **and** Figure S6).

In cluster 3-OUT, the trajectory of genes from *Pto AvrRpm1*-treated samples does not remarkably differ from mock treatment, with a pattern of steady expression throughout the course of the experiment and a mild increase of expression from 4 to 6 hpi (Figure 4a). Genes that comprise this cluster mainly fall into GO terms associated with the photosynthetic machinery (Figure 4b). These data indicate that photosynthesis at the OUT area of infection does not seem to be altered by pathogen infection as opposed to the IN area (Figure 3b-4b **and** Figure S5-S6) correlating with zonal photosynthesis efficiency values shown in Figure 1c and as previously reported (33).

### Novel zonal immune cell death transcriptional indicators can be elucidated from pairwise comparisons between time, treatment and area

In order to identify robust immune cell death markers that are exclusively upregulated at the site of cell death (IN area) we conducted a pipeline of differential expression analysis that consisted of concatenated pairwise comparisons considering the three variables in our experimental design: time, treatment and area (Figure 5a). Since the highest degree of differential expression between treatments took place at 4 and 6 hpi (Figure 2a), we carried out the comparisons at these two time points independently. Firstly, we focused on the time variable and selected genes that were confidently upregulated at the IN area of *Pto AvrRpm1*-infected plants, either at 4 and/or 6 hpi, compared to 0 hpi (1^st^ filter: FDR < 0.05 and log_2_FC>2). Secondly, we removed genes also upregulated at 4 and/or 6 hpi at the IN area in mock controls (2^nd^ filter: FDR < 0.05 and log_2_FC>2). Since we aimed to find genes only upregulated at the IN/cell death area, next, we removed the genes that were upregulated by bacterial inoculation at the OUT area at least to half of the levels than in the IN zone (3^rd^ filter: FDR <0.05 and log_2_FC<1). Finally, from the genes that met those three criteria, we kept those that were differentially upregulated at IN compared to the OUT area in *Pto AvrRpm1*-infected plants (4^th^ filter: FDR < 0.05 and log_2_FC > 2) (Figure 5a)

**Figure 5.**
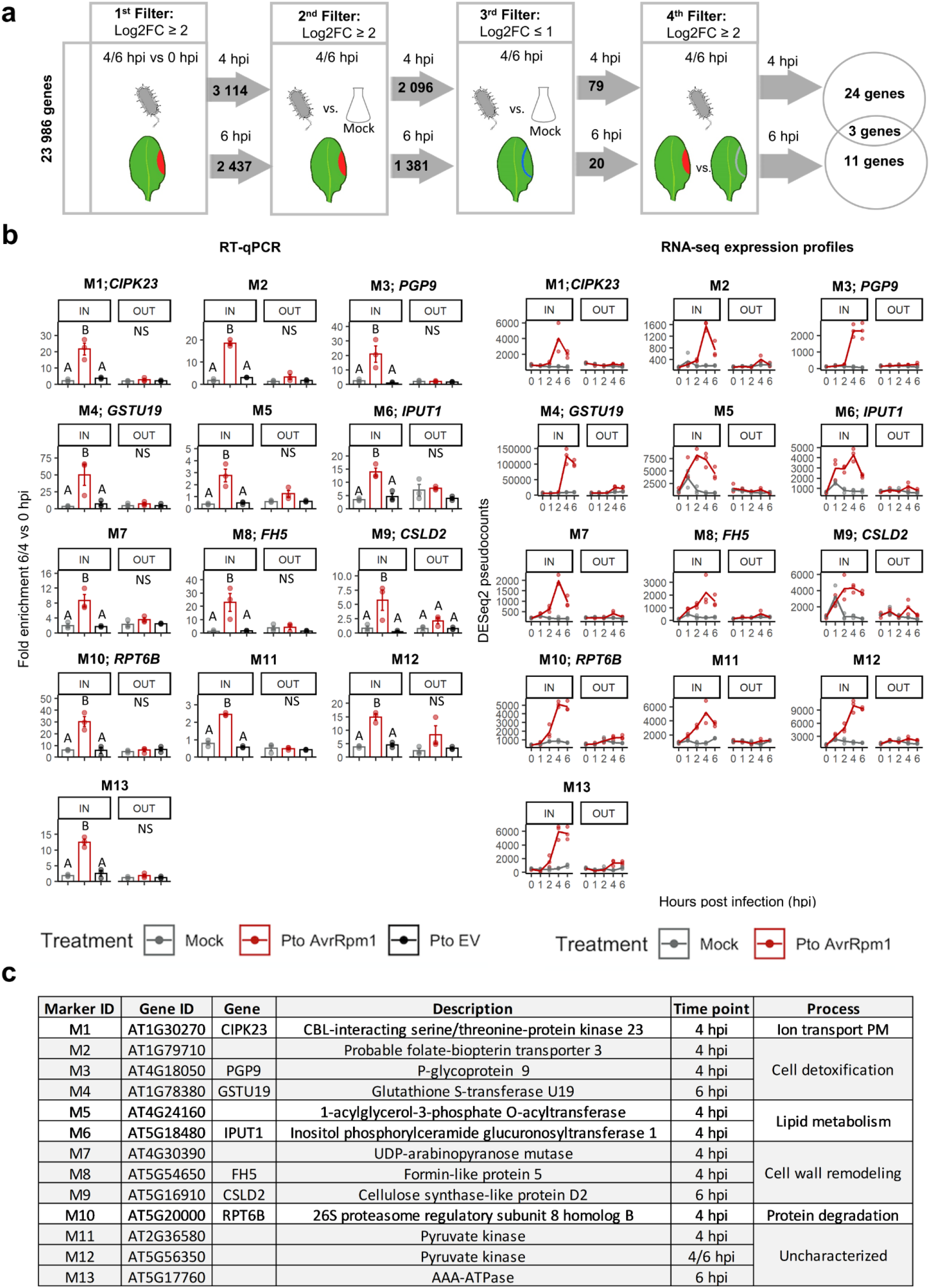
Identification of immune cell death markers specific for the IN area of infection. **(a)** Schematic representation of the sequence of filters applied to identify indicators. Four filters were concatenated considering the three variables of our experimental design: time, treatment and tissue area. Briefly, in the first filter, we selected genes differentially upregulated from 0 to 4/6 hpi (FDR < 0.05 and log_2_FC > 2) at the IN area (colored in red) upon bacterial infection. From the genes that passed this first filter, we selected those that were exclusively upregulated (FDR < 0.05 and log_2_FC > 2) due to bacterial infection at the IN area at 4/6 hpi. Subsequently, from the genes that made it into the third filter, we selected those that were not highly upregulated in the OUT area (colored in blue) upon bacterial infection at 4/6 hpi (FDR < 0.05 and log_2_FC < 1). Finally, we applied a fourth filter to discard genes that could potentially be basally upregulated at the OUT area upon pathogen treatment at 4/6 hpi (FDR < 0.05 and log_2_FC > 2). The starting number of genes and the genes passing the different filtering criteria are indicated. **(b)** RT-qPCR and RNA-seq expression profiles of marker genes that behave as *bona fide* immune cell death indicators. Relative expression levels to the housekeeping gene *EIF4a* were represented as fold enrichment between 4/6 and 0 hpi. Error bars represent standard error of the mean from three independent experiments. Letters indicate statistically significant differences between treatments following one-way ANOVA with Tukeýs HSD test (α = 0.05) performed independently at IN and OUT. NS (non-significant after one-way ANOVA). Exact p values are provided in **Table S2. (c)** List of HR indicators along with their gene ID, gene name and description.

A total of 31 genes passed all 4 filters, constituting a set of potential immune cell death indicators (Figure S7). From these, 24 were extracted from the 4 hpi dataset and 11 from the 6 hpi dataset and 3 from both time points (Figure S7). The expression profiles of this putative immune cell death indicators can be visualized as DESeq2 pseudo-counts as a function of time at both areas of infection in Figure S8. The expression patterns of these 31 genes at 0 and 4/6 hpi was validated by real time quantitative PCR (RT-qPCR) using newly obtained biological samples (Figure S9). To ensure that the potential markers were exclusively upregulated as part of the immune cell death response triggered by effector-mediated bacterial recognition and not as part of the defense responses triggered by disease-causing bacteria, we also included samples inoculated with *Pto* DC3000 EV (*Pto* EV), a strain that causes disease but does not trigger immune cell death in Arabidopsis Col-0. Among the 31 genes tested, a total of 13 (10 of them at 4 hpi, 4 at 6 hpi, with one at both time points), behaved as *bona fide* immune cell death indicators (Figure 5b-c), showing a distinctive upregulation specifically triggered at the IN area by an immune cell death-causing bacterium.

### The AAA ATPase *At5g17760* promoter specifically drives expression of GFP to the IN area of infection, constituting a robust transcriptional live marker of immune cell death

In order to generate much needed tools to extend our understanding of how immune cell death unfolds at the infection site and its surrounding tissues using live tissue, we generated stable transgenic Arabidopsis plants expressing green fluorescent protein (3xGFP) under the control of the promoters of each of the 13 identified putative immune cell death marker genes. A nuclear localization signal (NLS) was fused to GFP to concentrate the signal in the nucleus and facilitate detection, which enabled us to distinguish promoter-driven fluorescence from the auto-fluorescence derived from immune cell death (42).

The behavior of these transgenic reporter lines was assessed upon immune cell death activation by syringe infiltration of a restricted area of the leaf of adult plants with *Pto* expressing different secreted bacterial effectors that are recognized by various NLRs (Figure 6a). In addition to *Pto AvrRpm1*, we also analyzed the response of these plants to *Pto* expressing *AvrRpt2* (*Pto AvrRpt2)* or *AvrRps4* (*Pto AvrRps4)*, which induce immune cell death in Col-0 plants via the CNL RESISTANT TO P. SYRINGAE 2 (RPS2) and the TNL RPS4, respectively (43, 44). As controls, we included mock, *Pto* EV and a non-pathogenic mutant strain secreting no effectors (*Pto hrcC^-^*) (45). Among all reporter lines tested, plants expressing *pAT5G17760:NLS-3xGFP* showed the most cell-specific, robust and clear GFP signal in the nuclei of the leaf regions upon infection with *Pto AvrRpm1* (Figure 6b). Activation of *pAT5G17760* was limited to the syringe-infiltrated area and could not be detected in the surrounding tissues. The same pattern was observed after infiltration with *Pto AvrRpt2* or *Pto AvrRps4* (Figure 6b), which indicates that *pAT5G17760* robustly responds to pathogen-mediated activation of different classes of NLR receptors. Importantly, infiltration with the mock solution or with non-HR causing bacterial strains did not activate *pAT5G17760*. It is worth noting that for microscopy imaging experiments we used a lower bacterial inoculum (O.D_600_ 0.01) to mimic more natural infection conditions and to delay the onset of immune cell death and tissue collapse (Figure 6a), which was necessary for microscopic detection of GFP (Figure 6b-e). At higher inoculum, rapid accumulation of phenolic compounds at the site of infection results in extremely high auto-fluorescence levels that hamper imaging. Together, our observations indicate that *pAT5G17760* activity is spatially regulated and confined to the area undergoing immune cell death. Thus, the transgenic reporter line *pAT5G17760:NLS-3xGFP* constitutes a very useful tool to monitor this process *in planta*.

**Figure 6.**
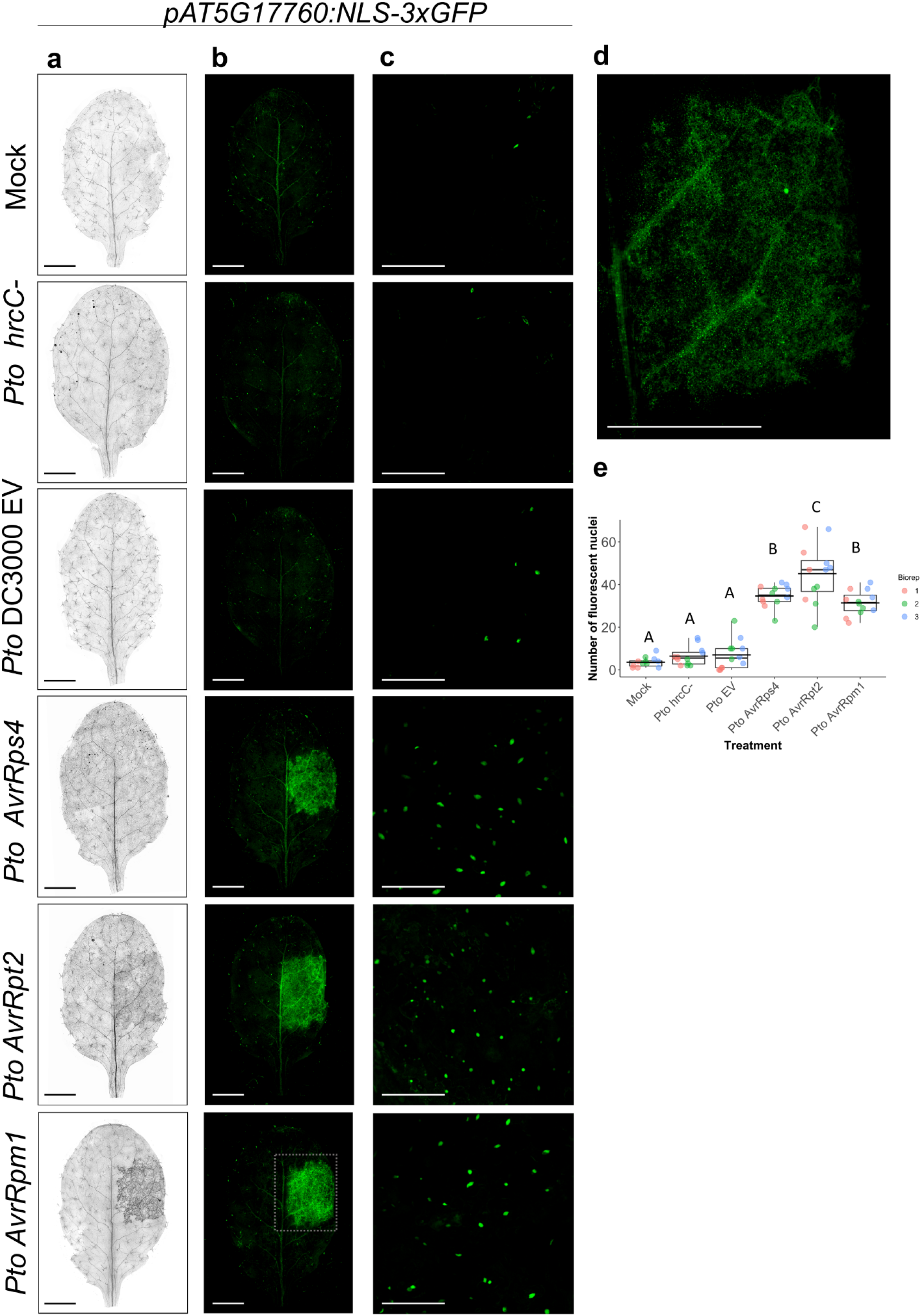
*AT5G17760* encodes an AAA-ATPase and is a reliable immune cell death indicator specifically induced at the IN area by activation of different classes of NLR receptors. **(a)** Representative images of trypan blue-stained leaves from *pAT5G17760:NLS-3xGFP* Arabidopsis transgenics. A small region of 4-week-old *pAT5G17760::NLS-3xGFP* leaves was syringe-infiltrated with *Pto* expressing the effectors *AvrRpm1*, *AvrRpt2* or *AvrRps4* at 1*10^7^ colony-forming units (CFU)/ml (O.D_600_ = 0.01). Besides mock treatment, the non-cell death-causing bacterial strains *Pto* DC3000 EV and *Pto* DC3000 *hrcC-* were included as negative controls. Images were taken 16 hpi. Scale bar 3 mm. **(b)** Representative fluorescent microscopy images from *pAT5G17760:NLS-3xGFP* Arabidopsis leaves infiltrated with the same pathogens and controls as in (a). Images were taken 16 hpi on a Leica DM6 microscope prior to trypan blue staining. Scale bar 3 mm. **(c)** Confocal microscopy images of the inoculated area as seen in **(b)**. Expression of p*AT5G17760* is detected as green dots corresponding to nuclei with positive *GFP* signal. Scale bar 100 µm. **(d)** Representative close-up image of a *Pto AvrRpm1*-infected leaf expressing *pAT5G17760:NLS-3xGFP* at 16 hpi. Scale bar 3 mm. **(e)** Quantification of fluorescent nuclei from confocal pictures in (c). Nuclei count was performed using ImageJ software. Data is representative of three independent experiments each one of them containing 4 leaves. Letters indicate statistically significant differences in number of nuclei following one-way ANOVA with Tukeýs HSD test (α = 0.05). Exact p values are provided in **Table S2**.

The *AT5G17760* gene encodes a putative AAA ATPase of unknown function. A knock-out mutant of this gene did not show any obvious immune cell death phenotype (Figure S11). The lack of phenotype could be due to functional redundancy/compensation, a very common masking phenomenon in plants. Future in-depth analysis of all immune cell death marker genes identified in this work, including combinatorial genetics, will contribute to a better understanding of immune cell death. This set of genes constitutes an invaluable tool to zonally discriminate cells undergoing pathogen-triggered cell death and mechanistically dissect this process.

## DISCUSSION

### Zonation of immune cell death in plants is underscored by distinct gene expression patterns and processes in dying vs by-stander cells

In plants, pathogen recognition via intracellular NLR receptors often results in an immune cell death reaction that helps preventing pathogen proliferation (46). This is a highly zonal response that takes place at the site of infection, whereby dying cells send signals to the surrounding tissues to activate defenses and block pathogen invasion. Traditionally, the plant immune system was considered strictly two-branched, with PTI being elicited by recognition of conserved pathogen patterns via cell surface receptors, and ETI recognizing pathogen effector proteins secreted into the plant cell via intracellular NLR receptors (2). Over the last decades, many efforts have been directed towards understanding the transcriptional reprogramming elicited during PTI and ETI (23, 31, 47–49). One of the major conclusions drawn from these studies is that whilst the repertoire of differentially expressed genes in the host is largely similar, ETI leads to a faster and more robust transcriptional response than PTI (5, 6, 23, 47, 50). These findings, together with emerging evidence showing additional levels of synergy and crosstalk between PTI and ETI has somewhat blurred the traditional PTI-ETI dichotomy (25, 51, 52). However, despite the large amount of time-resolved transcriptomic data produced (23, 31, 47, 53), the spatial consideration of immune cell death upon ETI activation has been partly overlooked, with only few studies pointing to its importance in regulating the process (27, 28, 54). It remains unclear whether and to what extent transcriptional reprogramming takes place at the vicinity of cell death compared to that occurring at the infected area upon bacterial infection.

Our experimental design (Figure 1a) considered the spatio-temporal angle of plant immune cell death to gain a better understanding of how this process is restricted to a few cells upon pathogen recognition and to define robust markers of the dying area over time. This is particularly important since in plants, cell death characterization has largely relied on biochemical and morphological hallmarks most of which are *post-mortem* and which in most cases do not provide unequivocal criteria (55, 56). We currently lack a set of genes that can be employed as gene indicators of cell death triggered by pathogens. *In silico* comparisons of transcriptome profiles at different developmental stages and upon environmental stresses leading to cell death, enabled identification of cell death indicators of developmentally regulated programmed cell death that can be used to detect or even isolate cells that are ready to die (7). The same approach did not lead to identification of reliable immune cell death markers, partly because the available datasets were not obtained on zonally resolved samples (7).

Here, differential expression analysis and clustering of genes based on expression profiles over time enabled us to infer biological processes taking place at each tissue area (IN/OUT) upon bacterial infection, giving us hints on how immune cell death can be spatially restricted. At the IN area, genes involved in a local immune response to ETI-triggering bacteria are greatly induced from 1 hpi onwards (Figure 2a **and** Figure S2). Accordingly, gene clustering of inoculated samples brings about an expression profile describing a pattern of upregulation at early time points (1 to 2 hpi) (cluster 1), and in which GO terms associated with an ETI response are enriched (Figure 3a-b). Tissue from the IN area, also contains a set of genes that cluster based on a pattern of steady expression followed by a peak of upregulation from 2 to 4 hpi (cluster 2). Genes belonging to this cluster are involved in diverse biological processes ranging from regulation of immunity, responses to JA and SA or protein turnover (Figure 3a-b). It is now well established that proteasome activity is strongly induced during bacterial infection and that certain subunits of the proteasome are required for efficient fine-tuning of immune responses in plants (57–59). Finally, we identified a strong transcriptional repression of photosynthetic genes at 4 hpi at the IN area (Figure 2b **and Figure S2b**), in accordance with the previously established notion that infection results in a global downregulation of genes associated with the photosynthetic machinery (60). Consistently, genes exhibiting an expression pattern of downregulation through time (cluster 3) are mainly involved in metabolic processes and certain aspects of photosynthesis (Figure 3a and 3b). This specific decrease in photosynthesis is particularly interesting in light of recent reports of the interplay between bacterial effectors and the chloroplasts, whereby certain effectors can suppress chloroplast functions and in turn, chloroplasts can adopt immune functions to fight off pathogens (61–63).

Our results also show that transcriptional reprogramming in host cells surrounding the infection area (OUT area) is less extensive with a lower number of differentially expressed genes than at the IN area, and starts later mostly from 4 hpi onwards. Remarkably, photosynthesis is not significantly affected at the OUT area, corroborating our *in vivo* measurements (Figure 1c) and previous findings (60). A relatively functional photosynthetic machinery may be key to maintain effective defense mechanisms and prevent these cells from dying as their neighbors. This finding might have been masked in previous transcriptional studies that have not taken into account the zonal nature of immune cell death, and reveals that the defense-growth trade-off may also have a marked spatial component that needs to be taken into account in future research. Besides photosynthesis, the OUT zone was characterized by a marked upregulation of wound/JA-related genes at 4 hpi (Figure 4a-b **and** Figure S2c). This response can also be observed at the IN zone but the level of upregulation at the OUT zone is remarkably higher (Figure S4), indicating an amplification in JA signaling at the cells surrounding the death zone. In addition, some of the JA-related genes are among those genes exclusively upregulated at OUT at 4/6 hpi, which indicates that they could potentially be used as zonal markers of the surrounding area (Figure 2b-c **and** Figure S4). It also indicates that not all JA components act at the same place and distance from the infection point. *In vivo* imaging of marker gene promoter activities of SA and JA signaling during ETI discerned two spatially distinct domains around the infection site, where JA signaling is thought to be important for regulating over-activation of SA signaling (29). Future studies that include mutants deficient in JA could provide mechanistic insights into how JA signaling contributes to the confinement of plant immune cell death. Our analysis also shows that some SA-signaling genes are among the upregulated IN-specific genes at late time points (Figure 2b-c **and Table S5**). Although originally considered antagonistic hormones required for immunity against pathogens with contrasting lifestyles (64), the interplay and synergism of these two phytohormones is now well established during ETI (35).

### Zonally resolved transcriptomic analysis of immune cell death in plants allows for the identification of robust biomarkers of the process

Robust biomarkers are essential to gain mechanistic knowledge of cell- or tissue-specific processes. In mammals, the extensive mechanistic knowledge of molecular constituents underlying regulated cell death has enabled the use of biomarkers for detection of tumor cells or aberrant cell death processes in cancer patients (65, 66). The field of immune cell death in plants is gaining momentum thanks to recent major discoveries that in one hand are leading to a redefinition of the PTI-ETI relationship and on the other, have provided mechanistic insight into how NLRs become activated and form supramolecular complexes that mediate cell death (5, 6, 13, 14, 25, 26). However, amidst this exciting scenario, the conceptual framework of immune cell death zonation is scarcely defined and will be key to understand its execution, spatial restriction mechanisms and define *bona fide* indicators of the process.

One of the main goals of our analysis was to define new markers of immune cell death. We made use of the RNA-seq data generated from IN and OUT areas in order to pinpoint gene indicators of immune cell death that can be used either as transcriptional markers or gene promoter markers for *in planta* detection of cells destined to die using live imaging. Applying stringent filters to our dataset we identified 13 genes that can be used as unequivocal transcriptional markers of zonally restricted cells that have activated a death program in response to pathogen perception via NLR activation (Figure 5c).

This marker set includes genes involved or putatively involved in various processes such as ion transport across the plasma membrane (M1), cell detoxification (M2, M3), lipid metabolism (M5, M6), cell wall remodeling (M7, M8, M9), protein degradation (M10), glycolysis (M11, M12), whereas one of these genes remains largely uncharacterized (M13) and encodes an AAA+ ATPase of unknown function. Interestingly, all these predicted functions are consistent with processes expectedly taking place on cells destined to die or that have started dying, although the function of most of these genes remains to be fully determined. This set of genes provide a glimpse of transcriptional regulation of immune cell death at the site of infection, the tip of the iceberg of the multi-level regulation of the process. For example, the fact that several genes are involved in cell wall remodeling highlights the importance of processes taking place in this extracellular compartment. In line with this, an increase of lignification at the edge of cells undergoing immune cell death was shown in the past and provided a clear picture of the zonal nature of this process (30). Interestingly, our transcriptome data clearly shows that many lignin biosynthetic genes are strongly and specifically upregulated at the IN zone at certain time points (Figure S10). How this cell wall lignification is regulated upon pathogen perception remains to be clarified and will be an interesting topic of research in the future.

Our data also reinforces the idea that the proteases involved in degradation of cell components during immune cell death are not particularly regulated at the transcriptional level. We observe specific upregulation of degradative processes at the IN zone such as autophagy, vacuolar degradation and proteasome-mediated processes and in fact, one of the marker genes is a proteasome subunit. However, we did not find any protease specifically upregulated at the IN zone, nor did any of them pass the filters to constitute a marker gene.

In parallel, the changes observed in marker genes involved either in ion transport across the plasma membrane or cell detoxification may be somewhat related with the predicted formation of a pore at the plasma membrane by pathogen-mediated activation of certain NLRs (16, 17). Although crucial pieces of this mechanism have been unveiled, knowledge is still scattered and we lack a more integrated picture that combines NLR activation with downstream processes, including cell death execution. In sum, our data provides a snapshot of how infected cells respond to pathogen recognition at the transcriptional level, compared to their neighbors, that are not directly exposed to the pathogen but respond to it. Importantly, this analysis has revealed a set of genes that are specifically upregulated at the IN zone and constitute robust markers of immune cell death, opening new paths to deepen our knowledge on the process.

Importantly, we present an Arabidopsis immune cell death reporter line stably expressing GFP under the control of the AAA+ ATPase *At5g17760* (M13), which shows extremely clear and strong expression exclusively at the inoculated area where pathogen recognition takes place via ETI, before the onset of cell death has become apparent (Figure 6). The other genes (M1-M12) constituted very clear qPCR markers but GFP promoter fusions did not result in a clear signal. This can be attributed to the limitations from defining an active promoter sequence or to the fact that their expression is not high enough to be detected via GFP, as microscopy is less sensitive than qPCR.

Interestingly, expression of the marker p*At5g17760:NLS-3xGFP* is similarly regulated by different classes of NLRs (CNLs and TNLs), revealing conservation of the process. Thus, this transgenic line constitutes a robust biomarker of immune cell death in plants triggered by activation of different NLRs that can be used for live monitoring of the process. Besides understanding the role of this gene in immune cell death, of particular interest will be to sort GFP-expressing cells of this transgenic line upon infection and adapt high-throughput cell death monitoring equipment used so far for animal cell death to describe and quantify the features and regulatory networks that define immune cell death in plants at a single-cell level.

## MATERIALS AND METHODS

### Plant and bacteria materials and growth

The *Arabidopsis thaliana* accession Col-0 was used for all experiments carried out in this study expect for electrolyte leakage. For electrolyte leakage, Col-0, *rpm1-3* (67) mutant of the NLR RPM1 and *at5g17760* mutant (GABI-KAT line 592F04_1) which carries T-DNA insertion in exon two, were used. Primers used for identifying the T-DNA mutant are listed in **Table S1**.

Seeds were sown on ½ Murashige and Skoog (MS) media supplemented with 1% sucrose and stratified at 4°C for two days. Plants were grown in a controlled chamber with a photoperiod of 9 h light and 15 h dark with white fluorescent lamps under 65% relative humidity. Seeds were germinated on plates and grown for 10-7 days, then individually transplanted to Jiffy pellets and grown for 3 additional weeks.

*Pseudomonas syringae* pathovar tomato *(Pto*) strains *Pto AvrRpm1*, *Pto AvrRpt2*, *Pto AvrRps4*, *Pto hrpC-* and *Pto* empty vector pVSP61 (EV) were grown on selective King’s B (KB) medium plates for 48 h at 28 °C. Bacteria was then resuspended in 10 mM MgCl_2_ and the OD_600_ adjusted to the appropriate inoculum.

### Bacterial inoculation and RNA-seq data collection

Bacteria were resuspended and the concentration was adjusted at 5*10^7^ colony-forming units or to an optical density measured at a wavelength of 600 nm (OD_600_) of 0.05. Fully expanded 7^th^ or 8^th^ rosette leaves were used for infiltration with either a mock solution (10 mM MgCl_2_) or *Pto AvrRpm1*. We syringe-infiltrated an area of roughly 3-4 mm at the side edge of leaves. Upon infiltration, the edge of the infiltrated area was underlined using India ink, and the total area infiltrated designated as “IN”. A 1 mm buffer zone next to the IN area was discarded and used as a reference to properly separate between the IN and the OUT zone, that expanded 1-2 mm towards the vein. Leaf tissue was separately collected from the IN and OUT area of infiltration at 5 different time points: 0, 1, 2, 4 and 6 hours by making use of a sterile scalpel. Leaf tissue was stored in 2 mL Eppendorf tubes and snapped-frozen in liquid nitrogen until the time of RNA extraction. Each sample collected consisted of tissue from six leaves derived from three different plants. For generation of three biological replicates from each condition (area, treatment and time), three independent experiments were performed. total sum of 60 samples −2 treatments (mock/infected), 5 time points (0, 1, 2, 4 and 6 hpi), 2 areas (IN/OUT) and 3 biological replicates-were used for RNA-sequencing.

For RNA library preparation, 1 μg of RNA from each sample was isolated using the NucleoSpin® RNA isolation kit (Macherey-Nagel, Hoerdt Cedex, France) following the manufacturer’s instructions. RNAseq was performed at the GeT-PlaGe core facility, INRA Toulouse. RNA-seq libraries have been prepared according to Illumina’s protocols using the Illumina TruSeq Stranded mRNA sample prep kit to analyze mRNA. Briefly, mRNA was selected using poly-T beads. Then, RNA was fragmented to generate double stranded cDNA and adaptors were ligated to be sequenced. 11 cycles of PCR were applied to amplify libraries. Library quality was assessed using a Fragment Analyzer and libraries were quantified by qPCR using the Kapa Library Quantification Kit (Kapa Biosystems, Inc, Wilmington, MA, USA). RNA-seq experiments have been performed on an Illumina HiSeq3000 using a paired-end read length of 2×150 bp with the Illumina HiSeq3000 sequencing kits.

### Read mapping and differential expression analysis

“FastQC” and “TrimGalore!” software was used for raw Illumina reads quality control analysis and trimming of reads containing adaptor- or vector-derived sequences, respectively (68). rRNA was detected and removed using “SortMeRNA 2.1b” software (69). Cleaned reads together with the transcriptome of *Arabidopsis thaliana* (as of 30 August 2018), including ncRNA, were used to quantify gene expression at transcript level using the software “Salmon v0.11.3” (70). Raw counts aggregated by gene were obtained using “tximport v1.14.2” and the result was used as input to “DESeq2” v1.26.0 (71, 72) to perform differential expression analysis. Then, genes adding up to less than 10 counts across all 60 samples were removed. The pre-filtered DESeq2 object contained 32,865 rows that turned to 23,986 after filtering. Counts normalized for sample size and regularized-logarithm transformed were used to produce PCAs.

Raw counts together with sample size information were used as input for DESeq2’s differential expression analysis. Simple pairwise comparisons based on a single factor were performed using DESeq2’s “result” function (73). while time course differential expression results were obtained using a likelihood ratio test as previously described (73). Genes with FDR below 0.05 and |log2FC| higher than 2 were considered as differentially expressed. FDR was calculated according to the Benjamini and Hochberg’s (BH) method (74).

### Gene clustering

Gene clustering was performed using Mfuzz v2.46.0 package under the R environment (39, 75) which is based on fuzzy c-means clustering algorithms. IN and OUT samples were independently analyzed. After time course differential expression analysis using DESeq2, only genes with an FDR <0.05 in the likelihood ratio test were selected for clustering.

The optimal number of non-overlapping clusters with a correlation value below 0.85 was 3 and 6 for *Pto AvrRpm1*-treated samples at the IN and OUT areas of infection, respectively. Subsequently, two highly redundant clusters were merged for OUT samples, yielding 5 final clusters. Genes that integrated each cluster derived from *Pto AvrRpm1*-treated samples were re-clustered for mock-treated samples in order to inspect the differences and similarities of trajectories between treatments over time. Between two and four mock-based sub-clusters were obtained for every infected-cluster. To avoid overlap, we reduced the number of sub-clusters to two in mock-treated samples. Each gene belonging to a cluster returned an associated membership score value (MSV) that ranged from 0 to 1 depending on how well it fitted the expression profile dictated by the overall genes comprising the cluster.

### Enriched Gene Ontology analysis

The set of genes that belonged to expression profile clusters or that exhibited differential expression were input into TAIR for Gene Ontology enrichment analysis for biological processes, which uses the PANTHER Classification system that contains up to date GO annotation data for Arabidopsis (76). The most specific term belonging to a particular family of GO terms was always selected for plotting. Only those GO terms exhibiting an FDR < 0.05 after Bonferroni Correction for multiple testing and a fold enrichment above 2 were selected for representation in dot plots.

### Identification of immune cell death indicators

For identification of immune cell death indicators, we concatenated four pairwise comparisons using DESeq2, in which we set different thresholds of log2FC, while keeping a stringent cut-off of FDR <0.05 throughout all comparisons. Briefly, we firstly selected genes that were upregulated (log2FC > 2) after *Pto AvrRpm1* infection at 4 or 6 hpi vs 0 hpi. From the genes that complied with this first filter, we selected those that were specifically upregulated in *Pto AvrRpm1*-infected vs mock-inoculated samples at 4 or 6 hpi (log2FC >2). From the genes that passed these two filters we kept those with a log2FC <1 at the OUT area in *Pto AvrRpm1*-infected vs mock-inoculated samples at 4 or 6 hpi. Since genes with log2FC near 0 do not usually have a low FDR, we kept our stringent FDR threshold while setting the log2FC threshold below 1 in order to capture with statistical confidence downregulated and only mildly upregulated genes at this tissue area. Finally, from the genes that met those three criteria, we kept those that were differentially upregulated at the IN area compared to the OUT area in *Pto AvrRpm1*-infected plants.

### Validation of gene expression by real time quantitative PCR

The same experimental setup used for RNA-seq data generation was followed for experimental validation by RT-qPCR including infections with *Pto AvrRpt2*, *Pto AvrRps4*, *Pto hrpC-* and *Pto* EV. Briefly, tissue was snap frozen and RNA isolated with the Maxwell® RSC Plant RNA kit (Promega). 1 µg of RNA was reverse transcribed into cDNA with the High-Capacity cDNA Reverse Transcription Kit with RNase inhibitor (Applied Biosystems^TM^). RT-qPCRs were performed with LightCycler® SYBRgreen I master (Roche) in a LightCycler® 480 System (Roche). Data was analyzed using the ΔΔCT method and represented as fold enrichment of the time point tested (4 or 6 hpi) relative to 0 hpi. Primers for RT-qPCR used in this study are listed in **Table S1**.

### Cell death analysis

Trypan blue staining of Arabidopsis leaves was performed by collecting whole leaves in 50 ml tubes (each leaf in a separate tube) at the specified time-points after treatment and covered with a 1:3 dilution of the stain. Tubes were incubated in previously boiled water for 15 min, and then cleared overnight with chloral hydrate on an orbital shaker. After removal of staining solution, leaves were covered in a 50% glycerol solution and photographed using a Leica DM6 microscope.

### Electrolyte leakage

Whole leaves from four to five-week-old *Arabidopsis* Col-0, *rpm1-3* or *at5g17760* (GABI-KAT: 592F04) grown in short-day with a photoperiod of 9h light and 15h dark, were infiltrated with *Pto AvrRpm1* at a wavelength of 600 nm (OD_600_) of 0.05 using a 1-ml needleless syringe. Leaf discs were dried and subsequently collected with a 0.8-cm-diameter cork borer from infiltrated leaves. Discs were washed in deionized water for 1 h before being floated on 2 ml deionized water (4 discs per biological replicate). Electrolyte leakage was measured as water conductivity with a pocket water quality meter (LAQUAtwin-EC-11; Horiba, Kioto, Japan) at the indicated time points.

### Chlorophyll fluorescence imaging

An IMAGING-PAM (Pulse-Amplitude-Modulated) M-Series Chlorophyll Fluorometer system (Heinz Walz, Effeltrich, Germany) was used to investigate spatio-temporal changes in photosynthetic parameters at the IN and OUT areas of infection (77). Plants were kept in the dark for 30 minutes before measurement. Plants were exposed to 2 Hz frequency measuring light pulses for Fo (minimum fluorescence in the dark-adapted state) determination. Saturating pulses (800 ms) of white light (2400 mmol photons.m-2 s-1) were applied for Fm (maximum fluorescence in the dark-adapted state) determination. The photosynthetic efficiency or maximum quantum yield of PSII photochemistry (Fv⁄Fm) was determined as (Fm-Fo)⁄Fm. The relative PSII electron transport rate (ETR) was calculated by performing a kinetic analysis for 10 minutes with 60 second pulses (78). Areas of interest (AOI) included IN and OUT in order to evaluate spatial heterogeneity. The measurements were taken after 0, 1, 2, 4 and 6 hpi. Results are shown from 6 different AOI.

### Generation of transgenic promoter reporter lines

A region of approximately −2.5 kb upstream of the transcription starting site of *AT5G17760* was amplified from Arabidopsis Col-0 genomic DNA by PCR and cloned into the pGGA (plasmid Green Gate A) entry vector to generate pGGA-pMarkerGene (79). Each entry vector was then recombined with the following plasmids: pGGB-SV40-NLS, pGGC-3xGFP, pGGD-RBCSt (D-F), pGGF-AlliYFP (seed coat selection cassette for transgenic seed selection) and pGGZ-empty destination vector. Primers used for cloning and sequencing the final constructs are listed in **Table S1**. All plasmids were transfected by electroporation into *Agrobacterium tumefaciens* GV3101 strain containing the plasmid pSoup and then transformed into Arabidopsis Col-0 by the floral dipping method (80). Transgenic seeds from transformed plants were identified as those displaying a clear fluorescence signal under the stereo microscope Olympus SZX18.

### Pathogen inoculation and microscopy of reporter lines

For microscopy of reporter lines, plants were grown as previously described. Leaves of Col-0 p*AT5G17760*:NLS-3xGFP were infiltrated in the IN area with either a mock solution (10 mM MgCl_2_) or different *Pto* strains. *Pto* strains expressing the following effectors were used: *AvrRpm1*, *AvrRpt2* and *AvrRps4*. As controls, the *Pto* EV and *Pto hrcC-* strains were also used. All *Pto* strains were infiltrated at a wavelength of 600 nm (OD_600_) of 0.01 for microscopy imaging. Leaves were imaged at 16 hpi. Whole leaves were photographed using a Leica DM6 microscope (Leica Microsystems) equipped with DFC365 FX 1.4 MP monochrome digital camera. Bright field and GFP filter pictures were taken of each leaf. Confocal images were obtained using a FV1000 Olympus confocal microscope with the following excitation/emission wavelengths for GFP: 488 nm/500 to 540 nm. Confocal microscopy images were taken of the epidermal layer (20 Z-stacks with stack size of 1 μm) and fluorescent nuclei were counted using ImageJ software.

## Supplementary Information

Supplementary information is available at Cell Death and Differentiation’s website

## Acknowledgements

The authors would like to thank Susana Rivas, who conceived and initiated the project, but declined to be author on the manuscript. Likewise, we thank Susana Rivas’s team for their help with the preliminary experiments and the plant tissue harvest for the RNA-Seq. We thank as well Sebastien Carrère from the Bioinformatics facility at the LIPM, for his bioinformatics preliminary analysis. We also thank Simon Stael (VIB) for helpful comments and inspiring discussions and all members from the Bacterial plant diseases and cell death lab for their insights and suggestions. We thank José Luis Riechman (CRAG) and Miguel Ángel Moreno-Risueño for help with the analysis. We would like to thank Kenichi Tsuda for sharing his RNA-seq data of previously published transcriptomic studies (23) and Ignacio Rubio-Somoza for providing us with the green gate plasmid pGGD-RBCSt (D-F).

## Author Contribution Statement

JS-L designed and performed experiments, analyzed and interpreted data and wrote the manuscript

IS designed and performed experiments and analyzed and interpreted data and helped writing the manuscript

NR-S designed and performed experiments, analyzed and interpreted data and helped writing the manuscript

MS performed experiments

UP performed experiments

VMG performed analysis and interpreted data

MB-F performed analysis and interpreted data

MV interpreted data and helped writing the manuscript

DR performed experiments, analyzed and interpreted data, and helped writing the manuscript.

NSC conceptualized the research, designed the experiments, interpreted data and wrote the manuscript.

## Ethics Statement

The present study did not require ethical approval.

## Data Availability Statement

All RNA-seq data generated during this study can be found at Short Read Archive SRP324081. All code used for analysis can be found at https://gitlab.com/molecular_data_analysis/ath_hypersensitive_response.

## LIST OF SUPPLEMENTARY TABLES

**TableS1.** Primers used in this study.

**TableS2.** Tukey HSD p-values obtained from statistical tests applied in the study.

**TableS3.** Lists of differentially expressed genes in Figure 2a.

**TableS4.** Lists of genes that are upregulated in Figure 2b.

**TableS5.** Lists of genes constituting each GO term in Figure 2c.

**TableS6.** Lists of genes comprising each cluster from *Pto AvrRpm1* and mock-treated. plants.

**TableS7.** Lists of genes constituting each GO term in Figure 3b.

---

**Figure S1.**
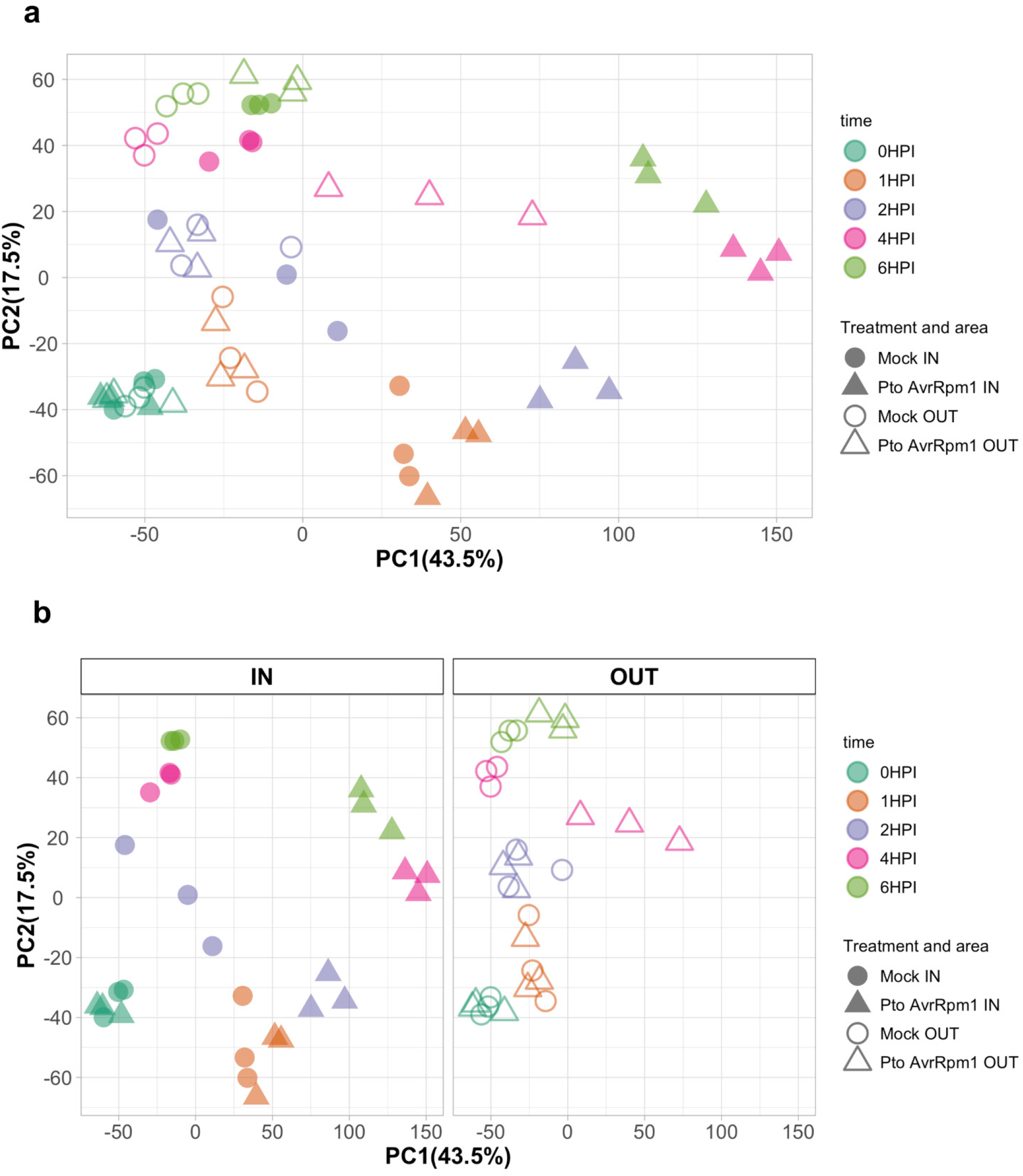
Principal component analysis (PCA) from the RNA seq-data. Circles represent mock-treated plants and triangles represent *Pto AvrRpm1*-infected plants. Different colors are assigned for each time point. (a) PCA comprising all data sets in our study (IN and OUT samples together). (b) PCA with IN and OUT data sets separated in order to ease visualization of the data.

**Figure S2.**
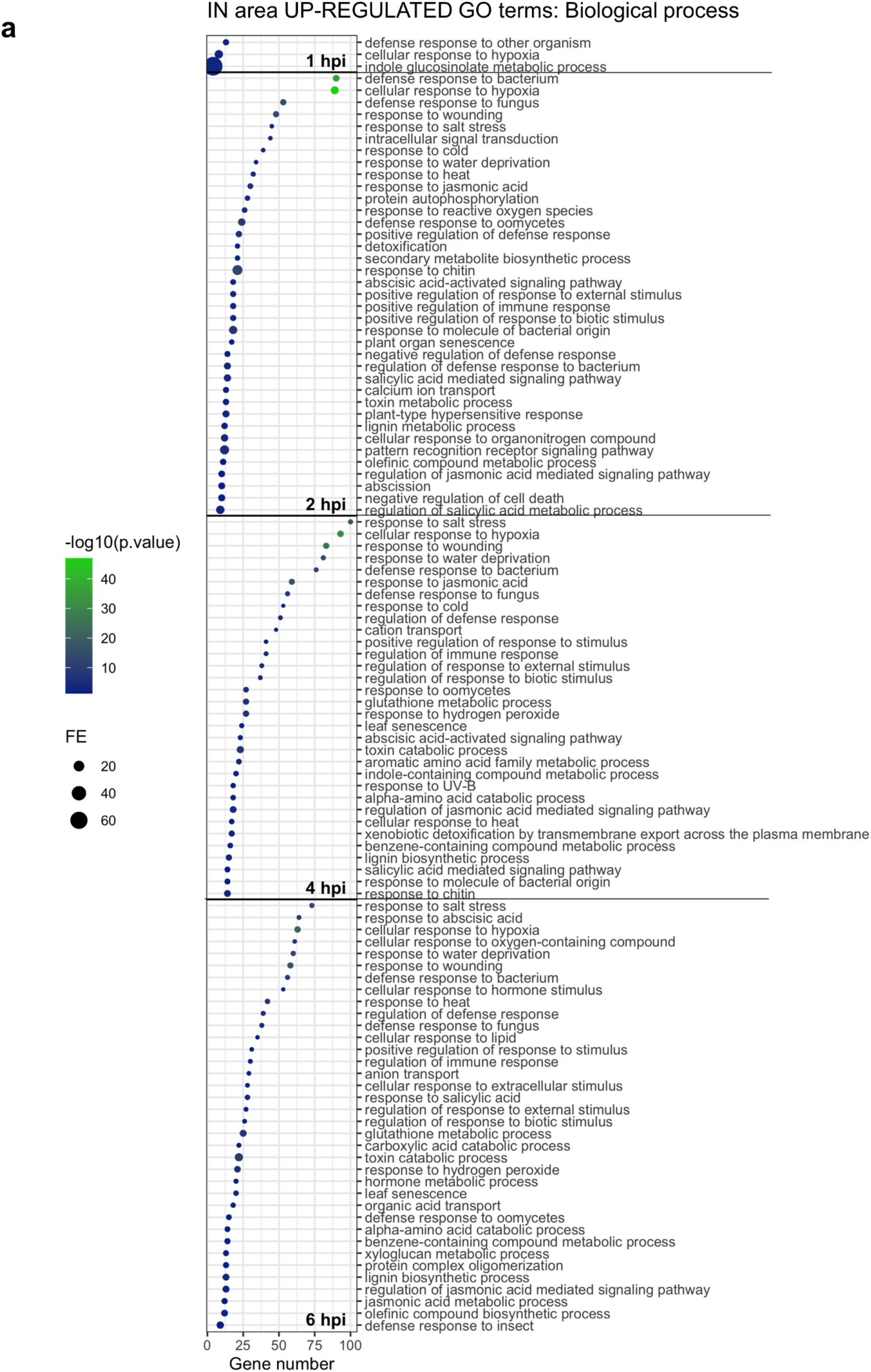
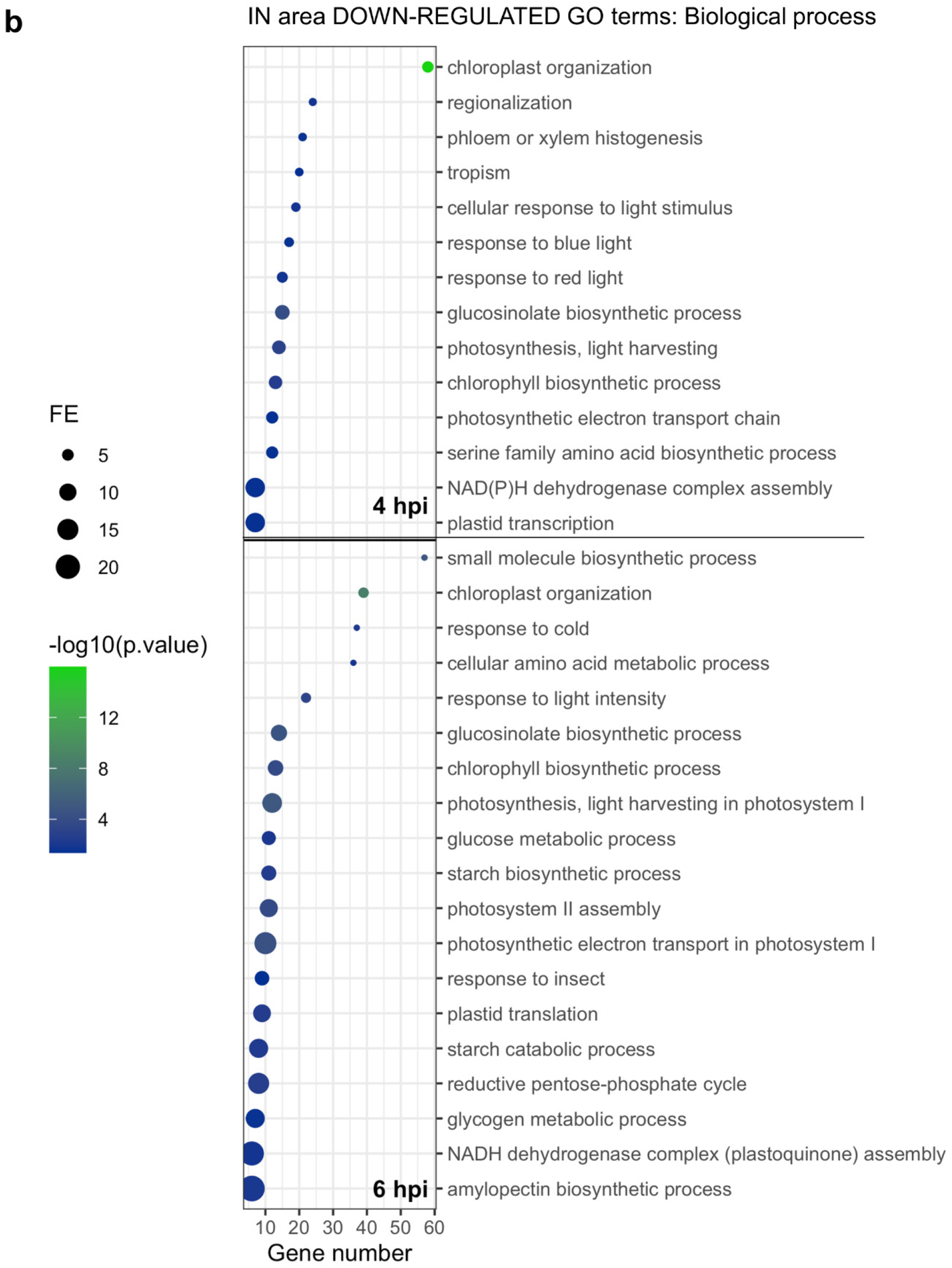
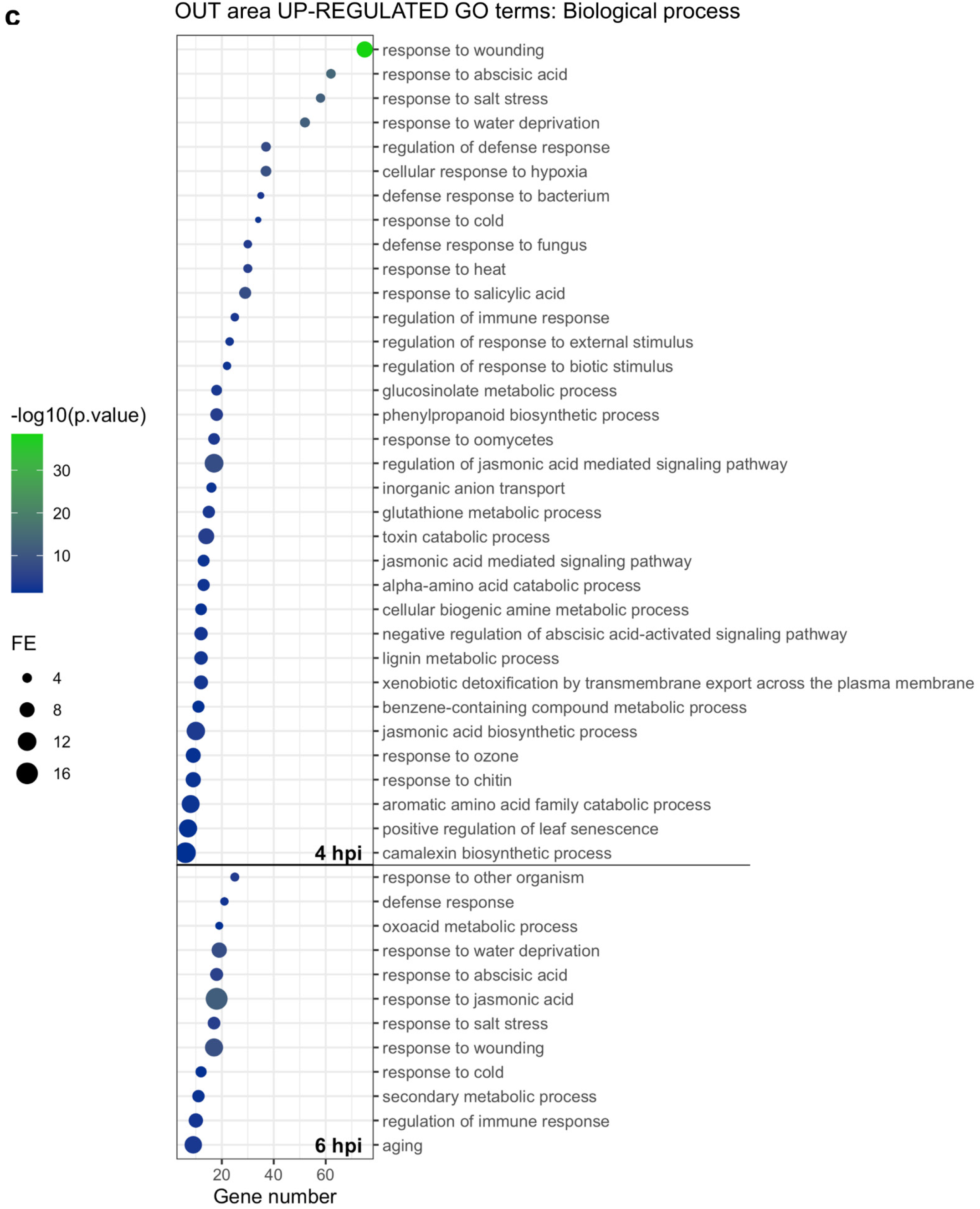
GO term enrichment analysis of upregulated and downregulated genes at each time after infection at the IN (a-b) and OUT (c) areas. The most specific term from each family term provided by PANTHER was plotted along with their corresponding gene number, fold enrichment and FDR (Bonferroni Correction for multiple testing) represented as log_10_. Only GO terms with a fold enrichment above 2 and FDR below 0.05 were plotted.

**Figure S3.**
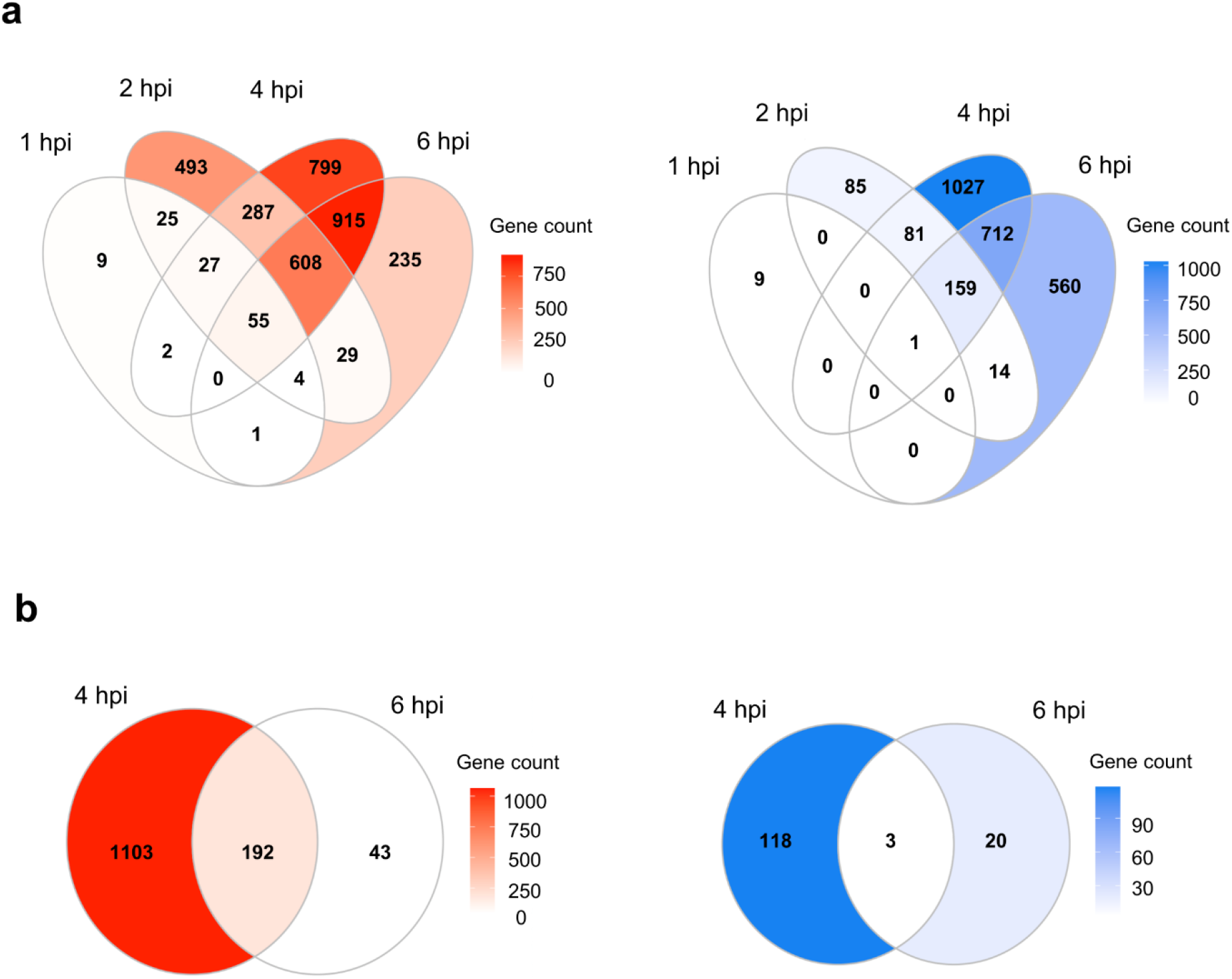
The majority of differentially expressed genes at both IN and OUT are specific to 4 and 6 hpi. Venn diagrams showing sizes of gene sets that are differentially expressed (red: upregulated and blue: downregulated) at IN **(a)** or OUT **(b)** at each time point.

**Figure S4.**
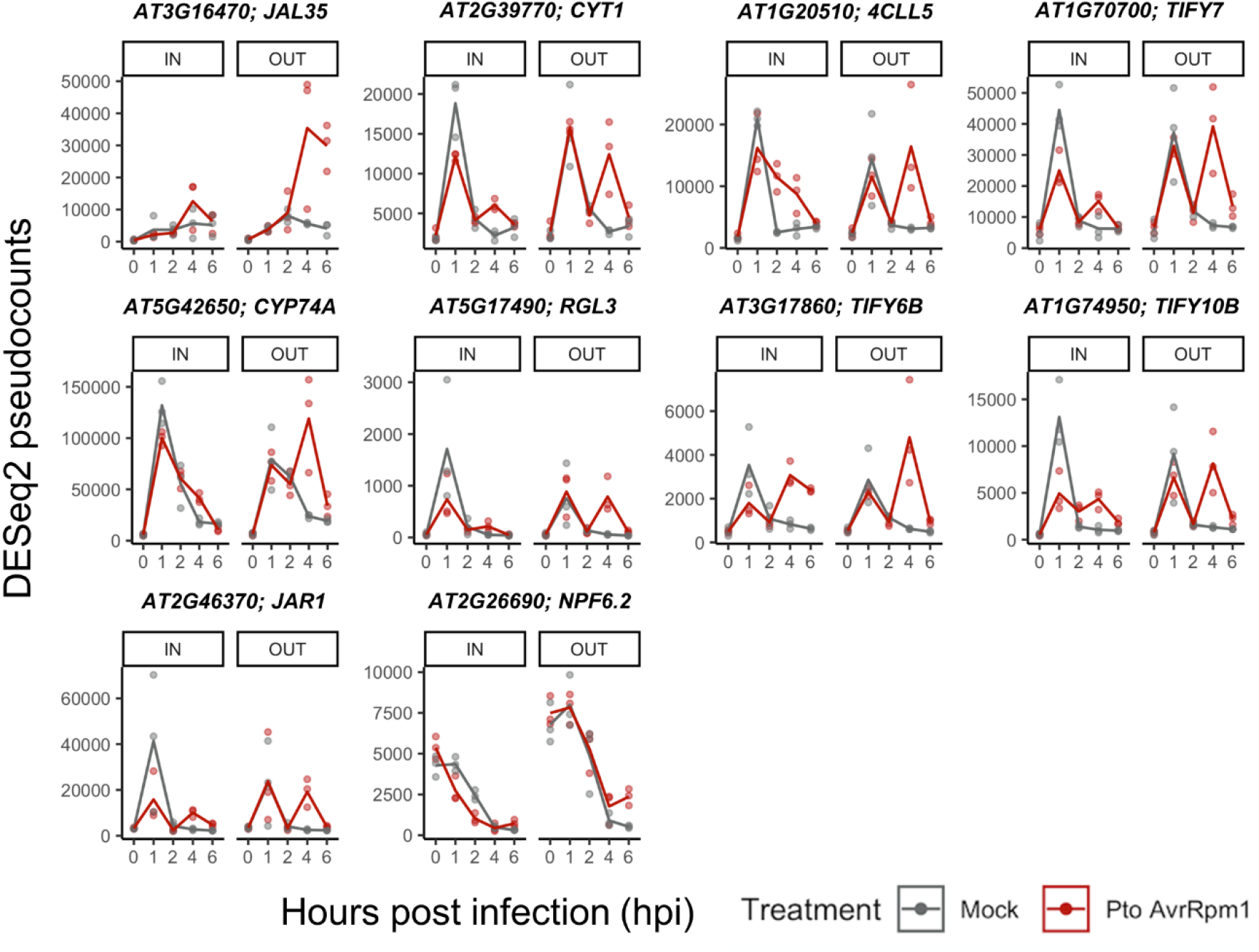
RNA-seq expression profiles of JA responsive genes exclusively upregulated at the OUT area upon *Pto AvrRpm1* infection. Gene expression of genes from *Pto-AvrRpm1* or mock-infected plants is represented as DESeq2 pseudocounts. *JAL35,* Jacalin-related lectin 35; *CYT1,* Mannose-1-phosphate guanylyltransferase 1; *4CLL5,* 4-coumarate--CoA ligase-like 5; *TIFY7,* Protein TIFY 7; *CYP74A,* Allene oxide synthase, chloroplastic; *RGL3,,* DELLA protein RGL3; *TIFY6B,* Protein TIFY 6B; *TIFY10B,* Protein TIFY 10B; *JAR1,* Jasmonoyl--L-amino acid synthetase JAR1; *NPF6.2,* Protein NRT1/ PTR FAMILY 6.2

**Figure S5.**
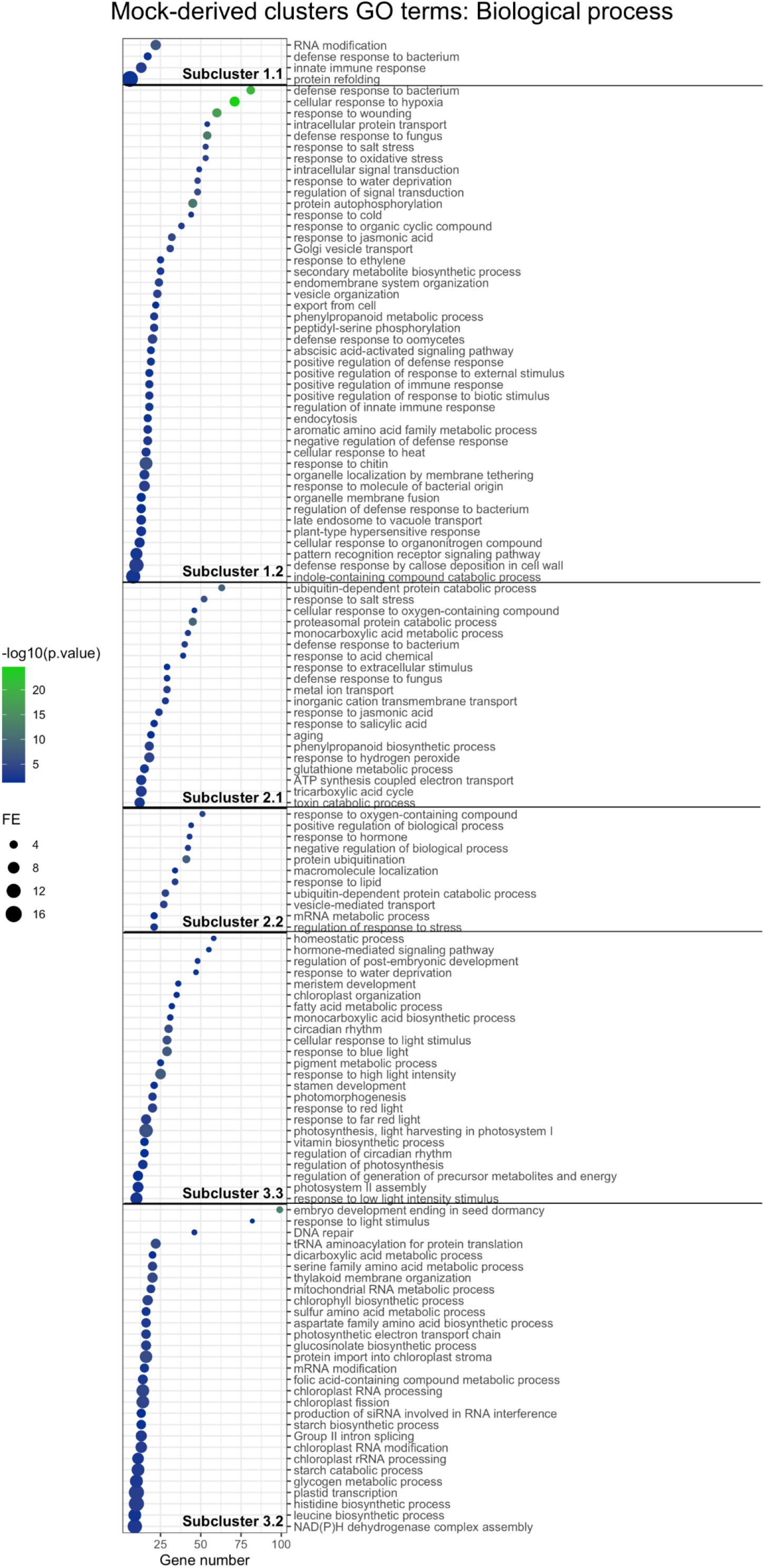
GO terms representing enriched biological processes derived from each sub-cluster in mock-treated plants at the IN area. From each cluster belonging to *Pto AvrRpm1*-treated samples, GO term enrichment analysis was performed on those genes that had a membership score value (MSV) above or equal to 0.7. The most specific term from each family term provided by PANTHER was plotted along with their corresponding gene number, fold enrichment and FDR (Bonferroni Correction for multiple testing) represented as log_10_. Only GO Terms with a fold enrichment above 2 and FDR below 0.05 were plotted. Sub-cluster 1.1 (638 genes; MSV >= 0.7 → 467 genes), sub-cluster 1.2 (2299 genes; MSV >= 0.7 → 1942 genes), sub-cluster 2.1 (2570 genes; MSV >= 0.7 → 1573 genes), sub-cluster 2.2 (1613 genes; MSV >= 0.7 → 649 genes), sub-cluster 3.1 (3172 genes; MSV >= 0.7 → 2391 genes), sub-cluster 3.2 (3256 genes; MSV >= 0.7 → 2557 genes).

**Figure S6.**
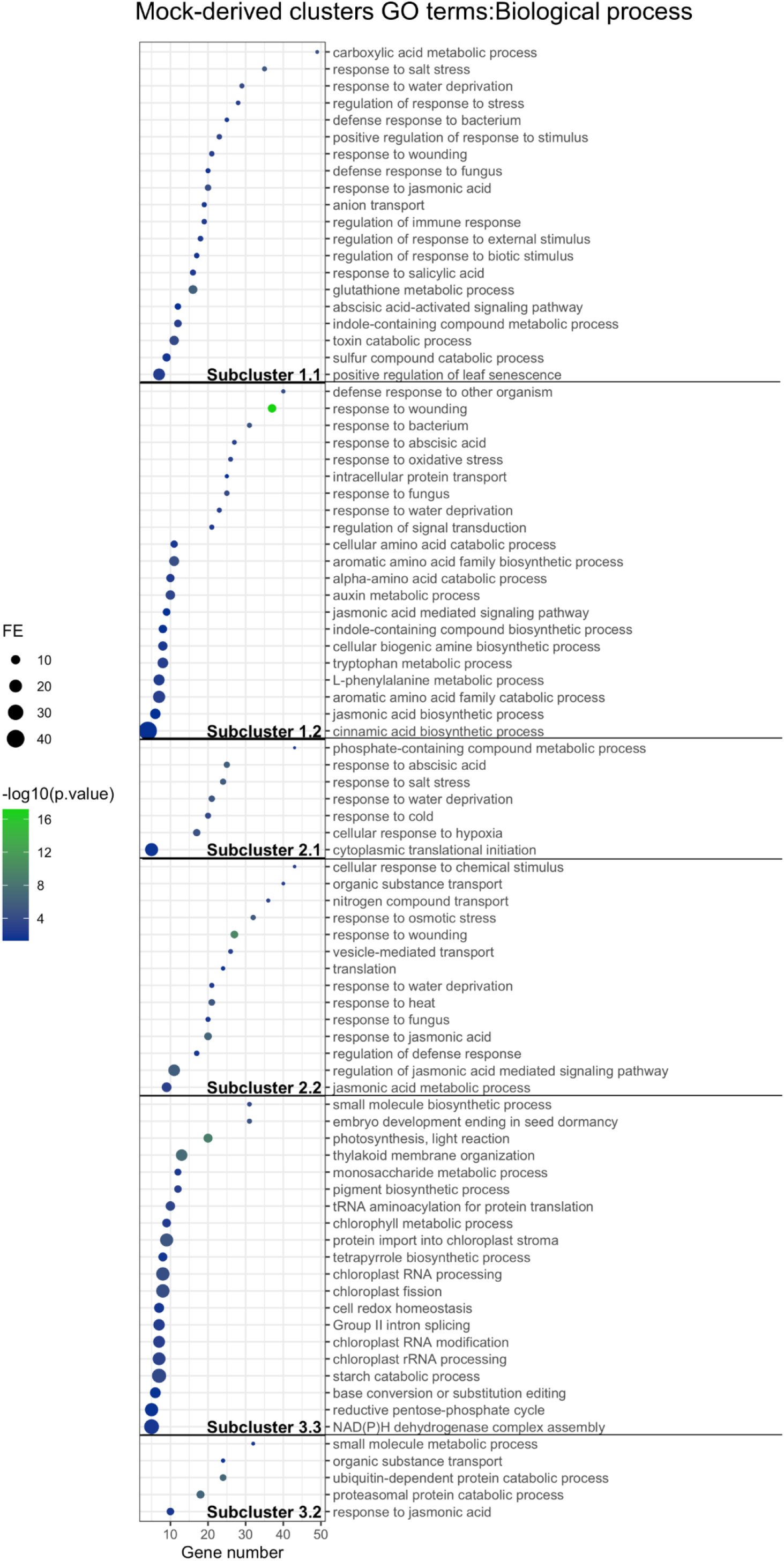
GO terms representing enriched biological processes derived from each sub-cluster in mock-treated plants at the OUT area. From each cluster belonging to *Pto AvrRpm1*-treated samples, GO term enrichment analysis was performed on those genes that had a membership score value (MSV) above or equal to 0.7. The most specific term from each family term provided by PANTHER was plotted along with their corresponding gene number, fold enrichment and FDR (Bonferroni Correction for multiple testing) represented as log_10_. Only GO Terms with a fold enrichment above 2 and FDR below 0.05 were plotted. Sub-cluster 1.1 (850 genes; MSV >= 0.7 → 319 genes), sub-cluster 1.2 (702 genes; MSV >= 0.7 → 183 genes), sub-cluster 2.1 (453 genes; MSV >= 0.7 → 286 genes), sub-cluster 2.2 (647 genes; MSV >= 0.7 → 389 genes), sub-cluster 3.1 (612 genes; MSV >= 0.7 → 555 genes), sub-cluster 3.2 (313 genes; MSV >= 0.7 → 257 genes).

**Figure S7.**
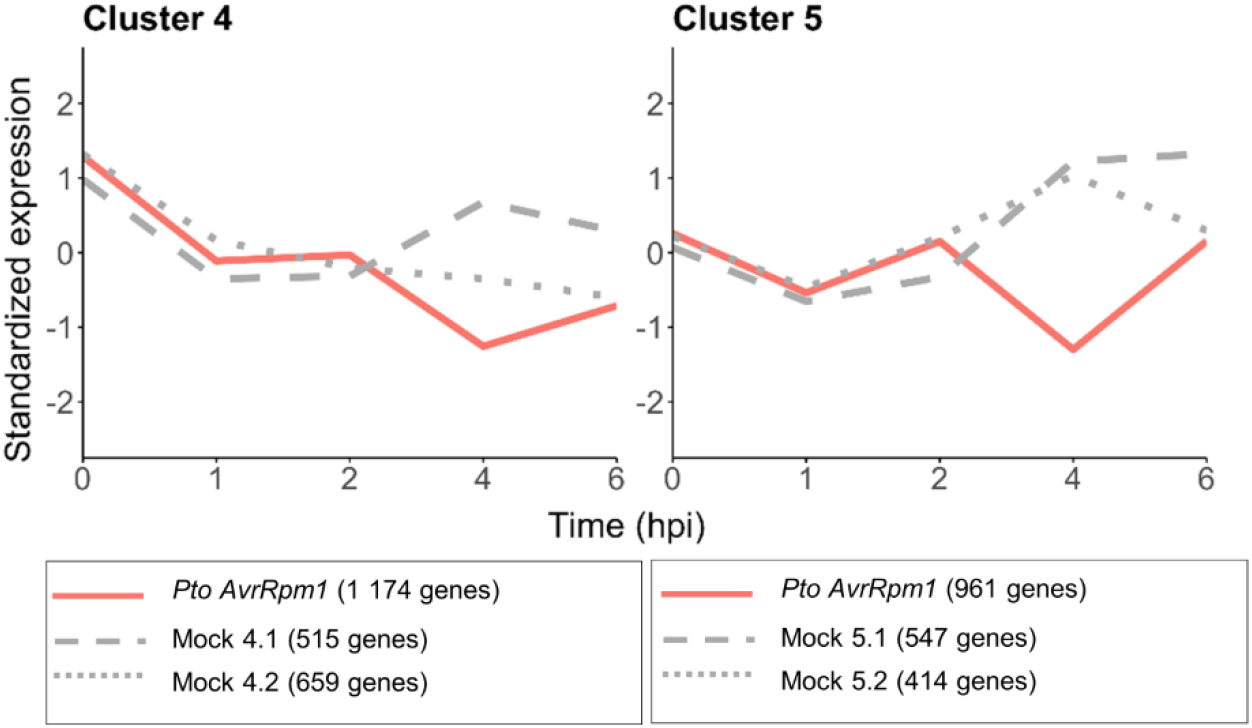
Clusters 4 (1,174 genes; MSV >= 0.7 → 57 genes) and 5 (961 genes; MSV >= 0.7 → 314 genes) from *Pto AvrRpm1*-treated plants at the OUT area share similar expression profiles and do not contain any relevant enriched GO terms associated with biological processes, possibly due to low gene number.

**Figure S8.**
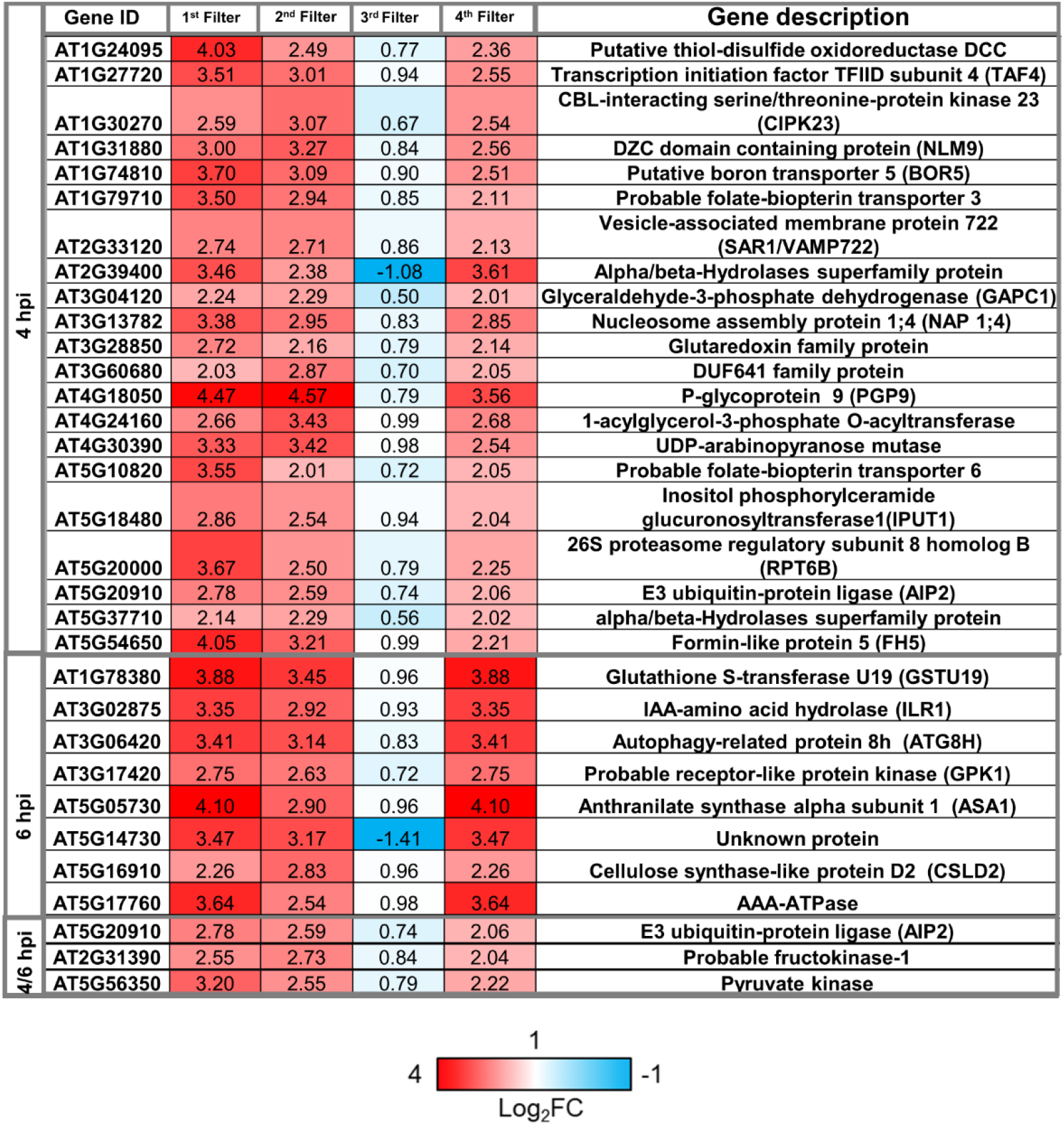
List of *in silico* HR indicators obtained after filtering at 4 and 6 hpi. Log_2_FCs from the the 1^st^, 2^nd,^ 3^rd^ and 4^th^ filters applied are indicated for each gene along with its corresponding gene description.

**Figure S9.**
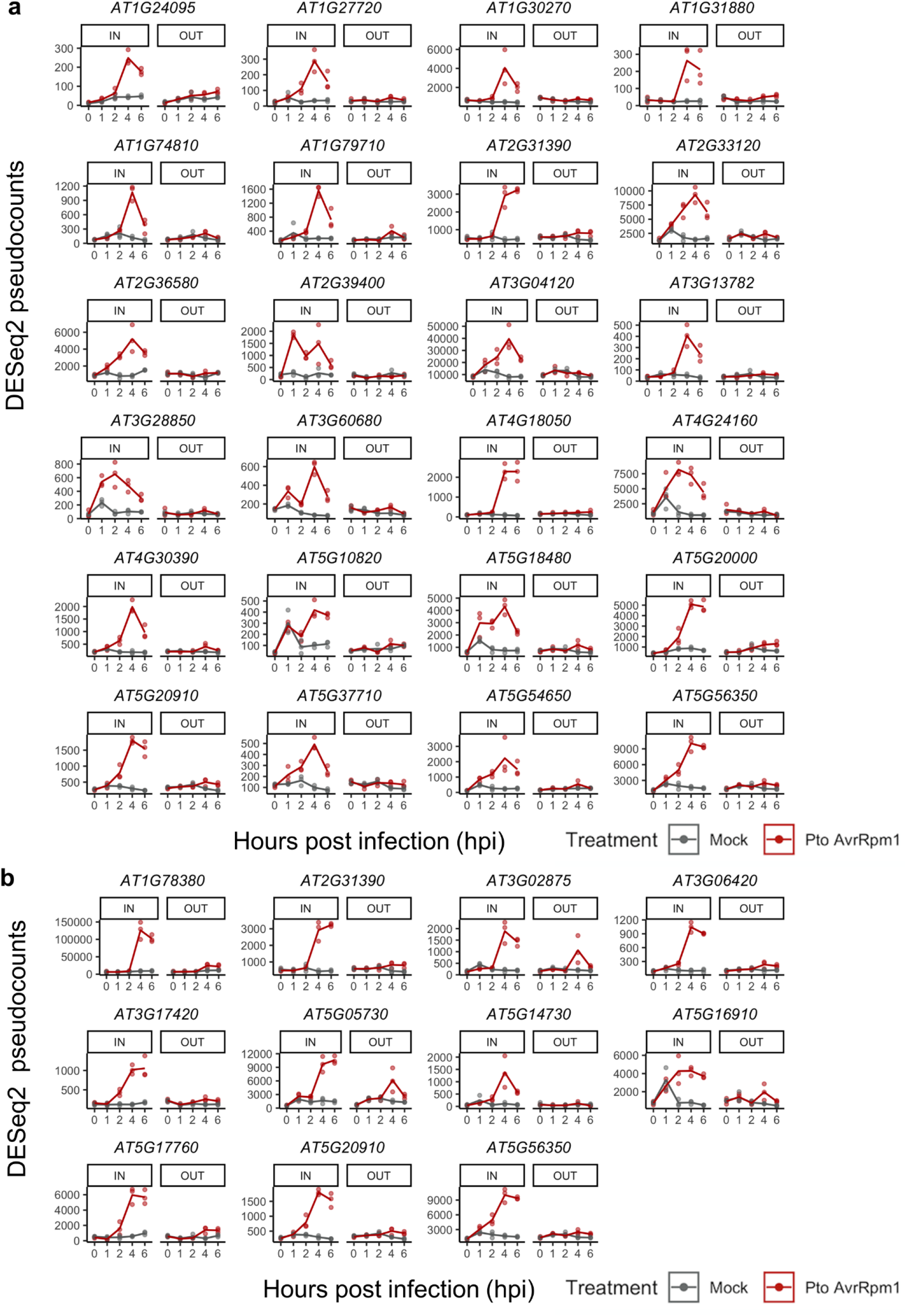
RNA-seq expression profiles of 4 (A) and 6 (B) hour candidate HR indicators at the IN and OUT areas of infection. Gene expression of genes from *Pto-AvrRpm1* or mock-infected plants is represented as DESeq2 pseudocounts.

**Figure S10.**
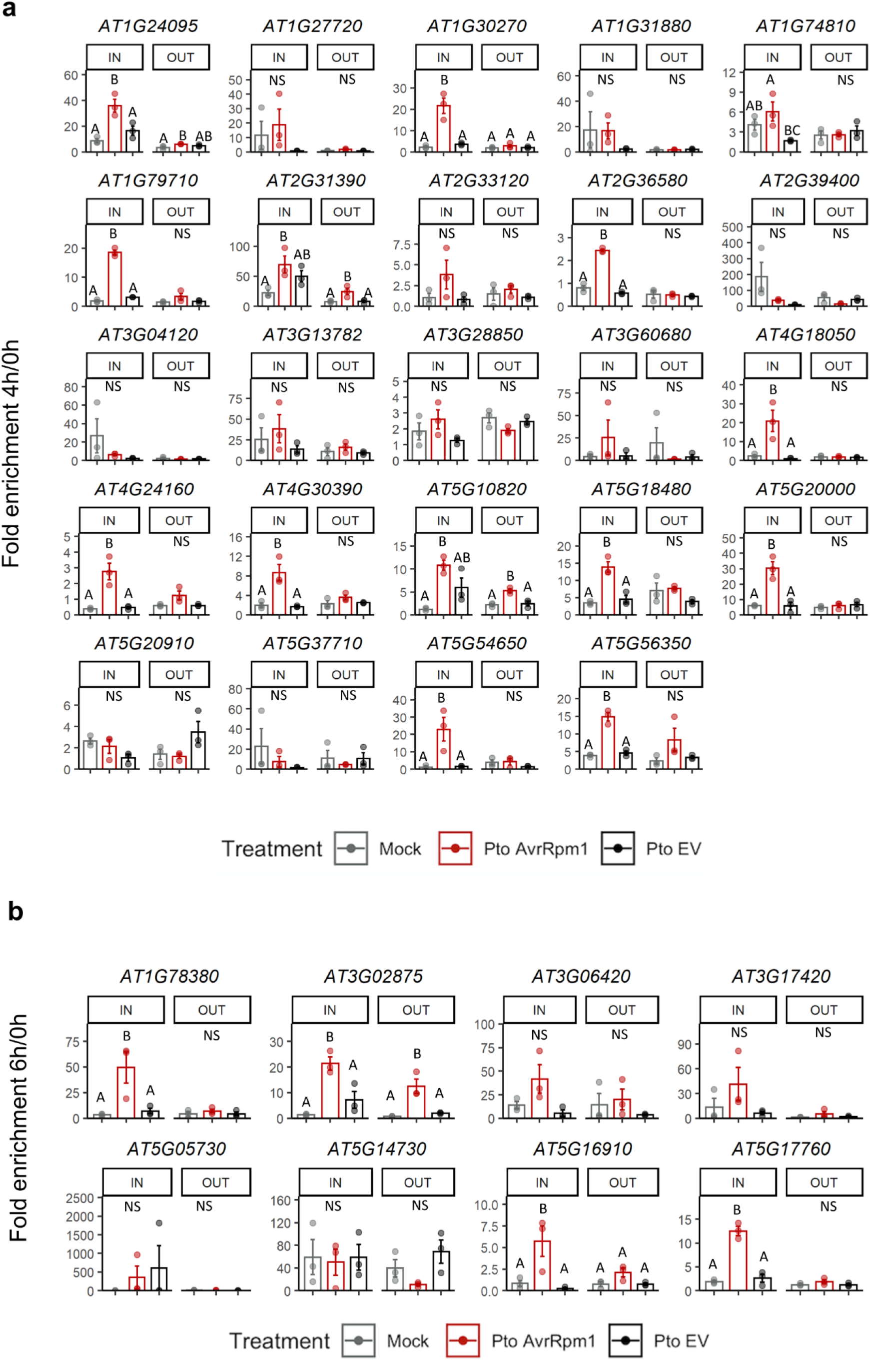
RT-qPCR of 4 and 6 hour transcriptional HR indicators at IN and OUT areas upon treatment with either mock, *Pto AvrRpm1* or *Pto* DC3000 EV. Relative expression levels to the housekeeping gene *EIF4a* were represented as fold enrichment between 4 **(a)** or 6 **(b)** and 0 hpi. Error bars represent standard error of the mean from three independent experiments. Letters indicate statistically significant differences between treatments following one-way ANOVA with Tukeýs HSD test (α = 0.05) performed independently at IN and OUT. NS (non-significant after one-way ANOVA). Exact p values are provided in Table S2.

**Figure S11.**
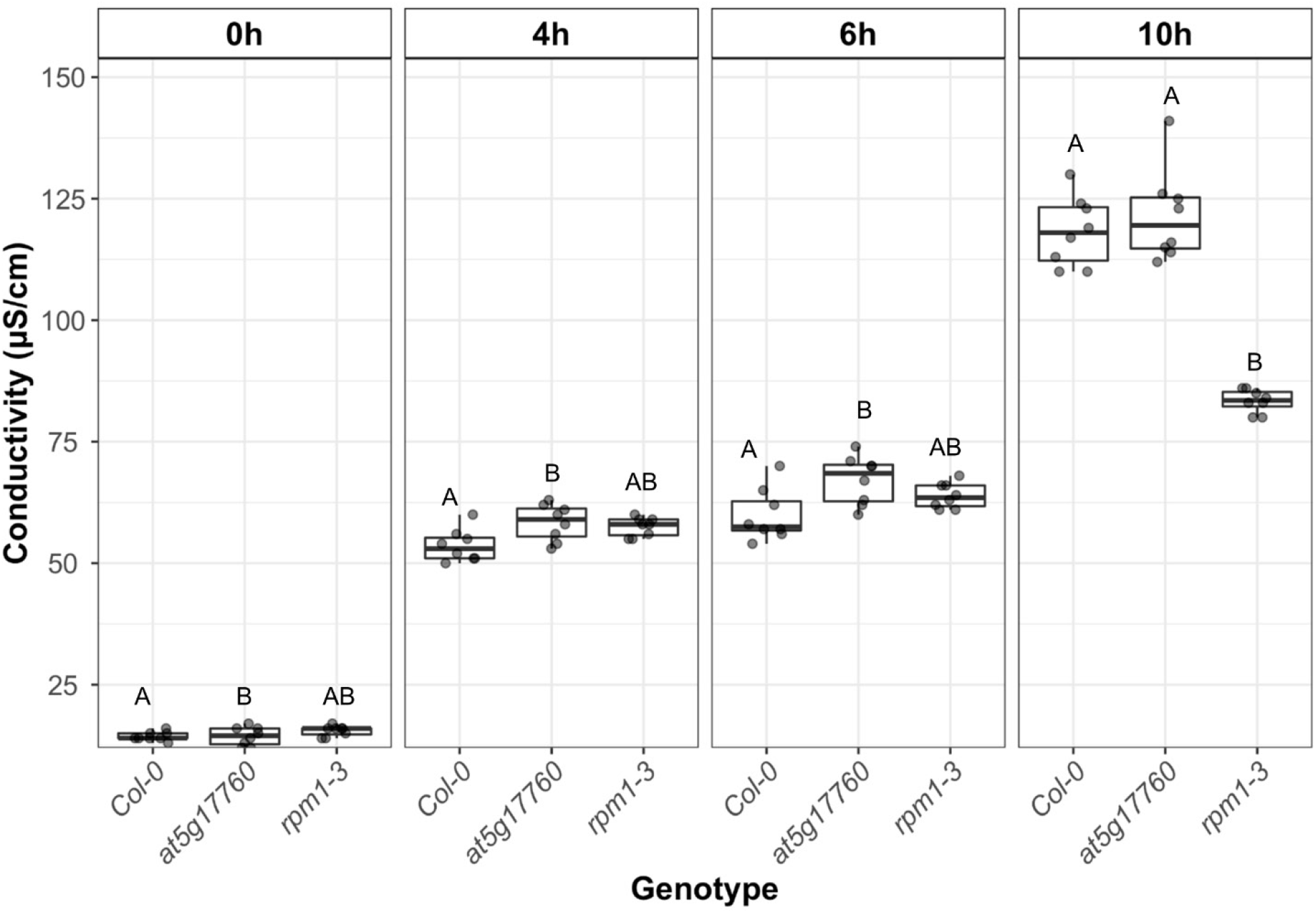
Onset of cell death in not compromised in an Arabidopsis mutant lacking AT5G17760. Four to 5 weeks-old plants were syringe-infiltrated with *Pto* DC3000 *AvrRpm1 at* 2.5×10^7 CFUs/O.D_600_=0.05. Conductivity measurements of electrolyte leakage from dying cells were recorded at 0, 4, 6 and 10 hpi. Dots represent data from 8 biological replicates consisting of 4 leaf discs each. Letters indicate statistically significant differences between genotypes following one-way ANOVA with Tukeýs HSD test performed at each time point. Exact p values are provided in Table S2.

**Figure S12.**
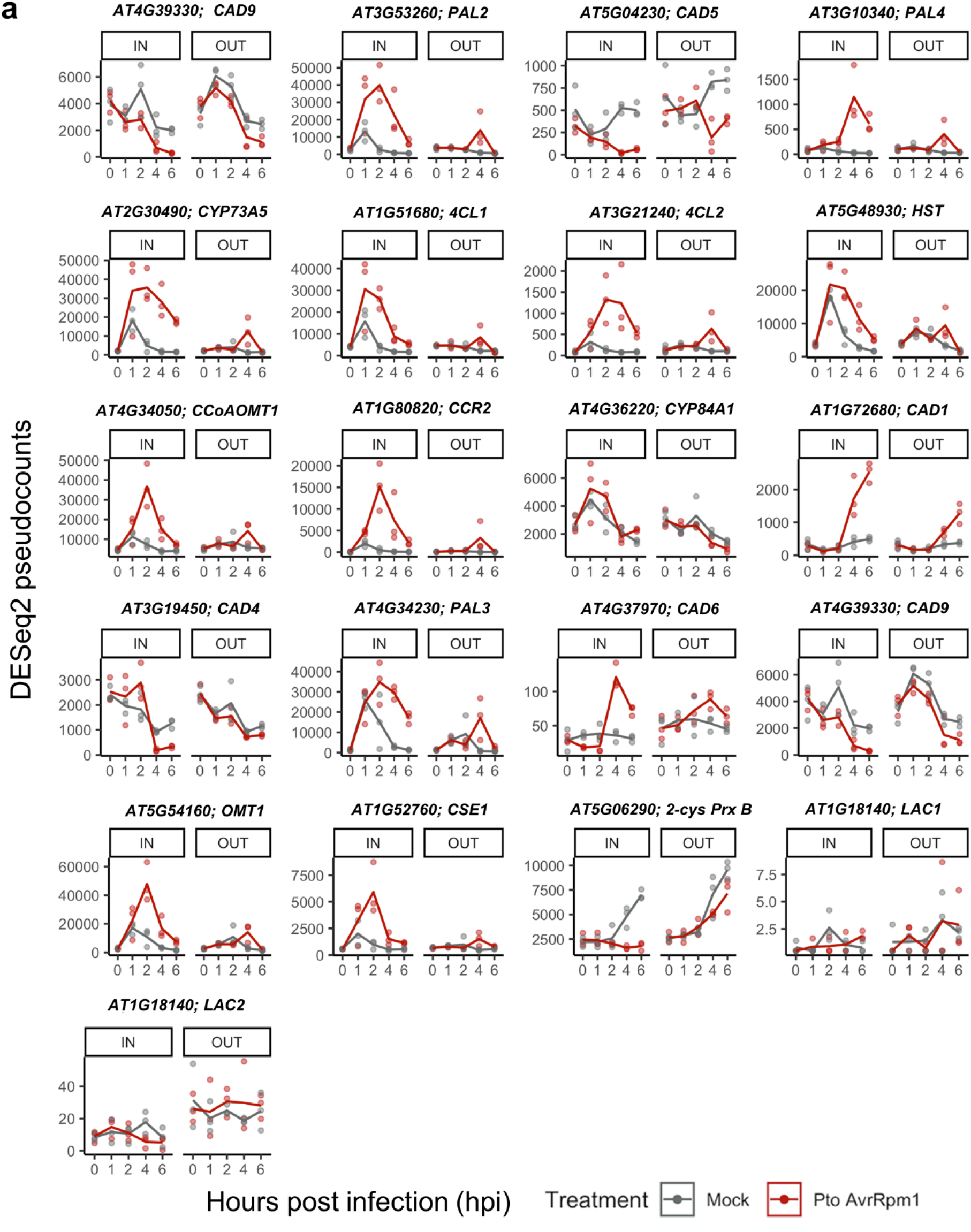
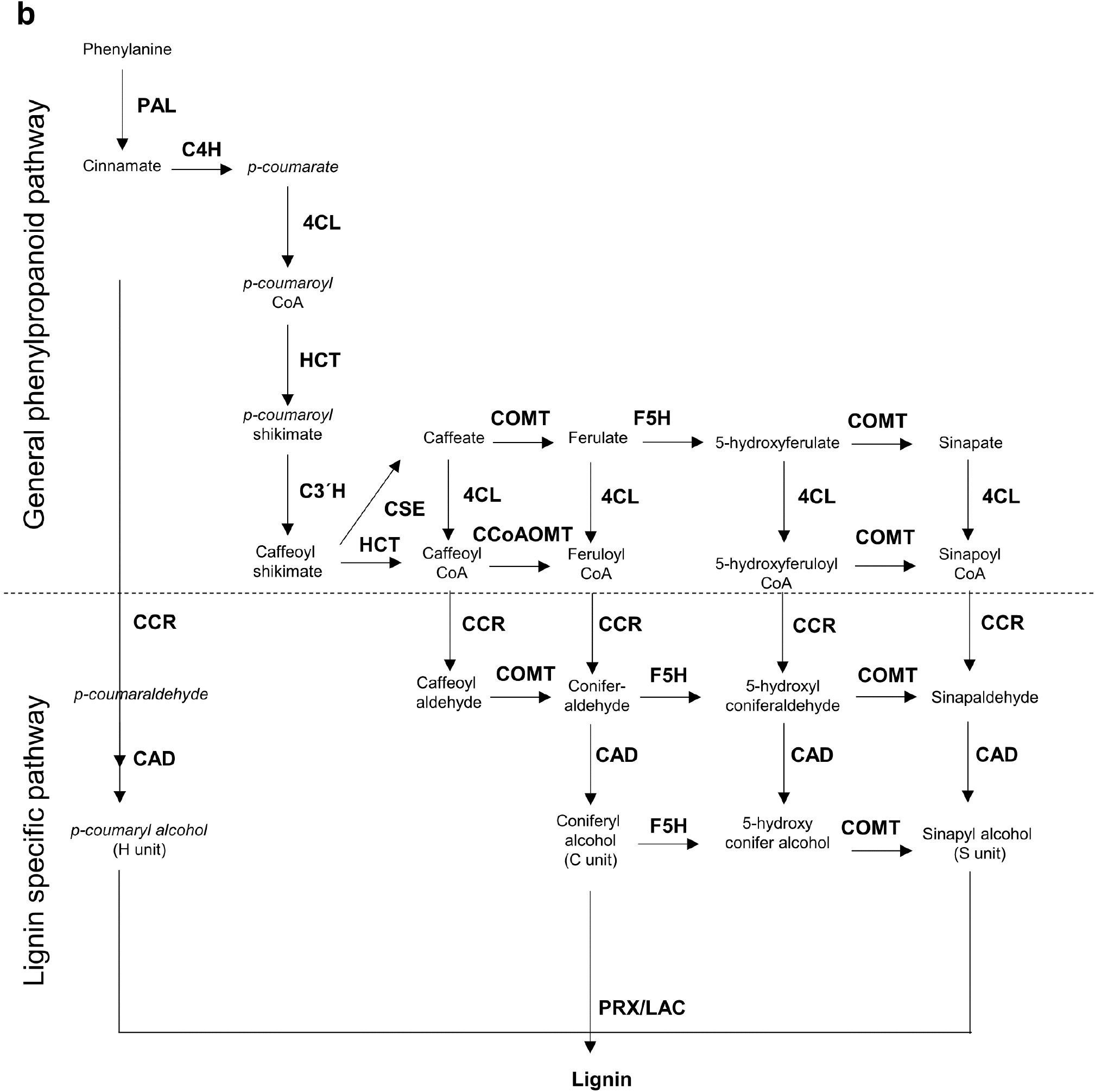
RNA-seq expression profiles of genes involved in lignin biosynthesis. **(a)** Gene expression of genes from *Pto-AvrRpm1* or mock-infected plants is represented as DESeq2 pseudocounts. **(b)** Scheme of lignin biosynthesis in plants. Black arrow indicates the canonical lignin biosynthesis in plants. Bold font indicates enzymes involved in the different steps of the pathway. PAL, phenylalanine ammonia-lyase; C4H, cinnamate 4-hydroxylase; 4CL, 4-coumarate: CoA ligase; HCT, quinateshikimate *p*-hydroxycinnamoyltransferase; C3′H, *p*-coumaroylshikimate 3′-hydroxylase; CCoAOMT, caffeoyl-CoA *O*-methyltransferase; CCR, cinnamoyl-CoAreductase; F5H, ferulate 5-hydroxylase; CAD, cinnamyl alcohol dehydrogenase; COMT, caffeic acid *O*-methyltransferase; CSE, caffeoyl shikimate esterase; PRX, peroxidase; LAC, laccase (Adapted from Meng Chie et al., 2018).

